# Molecular Shape Evolution of Cyanobacterial Circadian Clock Protein KaiC

**DOI:** 10.64898/2026.04.01.715469

**Authors:** Yoshihiko Furuike, Shuji Akiyama

**Author notes:** Shuji Akiyama, **Email:** (S.A.). Yoshihiko Furuike, **Email:** (Y.F.). **Author Contributions:** S.A. and Y.F. designed the research. Y.F. conducted sample preparations, biochemical experiments, and structure predictions. S.A. collected and analyzed small-angle x-ray scattering data. Y.F. and S.A. wrote the final version of the paper. **Competing Interest Statement:** The authors declare no competing interests.

## Abstract

A primitive form of clock protein KaiC has diverged into autonomous or passive time-measuring system in prokaryotes under selective pressures of day–night environmental changes caused by the rotation of Earth. However, the timing of such functional diversification and its structural basis remain unknown. Here we traced molecular shape evolution of older KaiCs by using X-ray solution scattering and structure prediction techniques. The result shows that the oldest ancestral KaiC emerged approximately 3.1 billion years ago as a moderately expanded and asymmetric double-ring hexamer, and subsequently evolved over a period of approximately 1 billion years into a compact and symmetric hexamer that is essential for achieving the self-sustained rhythmicity in extant cyanobacteria. In parallel with this compactification, the oldest KaiC branched into an oligomer composed of two hexamers approximately 0.5 billion years after its emergence. This is the direct experimental result demonstrating the early appearance of the prototypical dodecamer predicted by Kern and colleagues. It appears that this prototypical dodecamer gained a higher enzymatic activity during the next 0.4 billion years or so, and was passed down to non-cyanobacterial lineages as the passive timer capable of responding rapidly to environmental cues. Consequently, geological fluctuations over approximately 1 billion years since the earliest KaiC appeared caused the molecular shape of ancient KaiCs to evolve dramatically along the two distinct pathways.

**Significance Statement:** Molecular shape analysis has shed light on the 3-billion-year evolutionary history of clock protein KaiCs. The earliest KaiC emerged as a less compact and asymmetric hexamer, and later diverged along two distinct pathways. One led to the formation of a compact and symmetric hexamer as the core component of the self-sustained and temperature-compensated circadian oscillator found in modern cyanobacteria. The other reached the oligomerization of two hexamers as a prototype of the environmentally responsive timer found in non-cyanobacterial species. In the billion years following the birth of the oldest KaiC, ancient KaiCs adapted environmental alterations by dramatically evolving their molecular shape along the two directions.

## Introduction

Evolution is destined by the frequency of mutations and the strength of selective pressures (1-3). Gene mutations directly affect the function and structure of resulting proteins, and only organisms that have acquired resistance to environmental changes, drugs, or other stresses are selected. Under strong selective pressures, the adaptive evolution tends to accelerate in parallel with elimination, whereas under weak selective pressure, phenotypic changes occur gradually.

Circadian clocks are products of the adaptive evolution to day–night environmental cycles (4-13), exhibiting a self-sustained 24-hour rhythm, temperature compensation of the cycle length, and synchronization (14). The oldest timing system is considered to be ancestral forms of *kai* genes and their translational products, Kai proteins, found in extant cyanobacteria, non-cyanobacterial bacteria, and archaea (6, 11-13, 15) (Fig. 1A).

**Figure 1.**
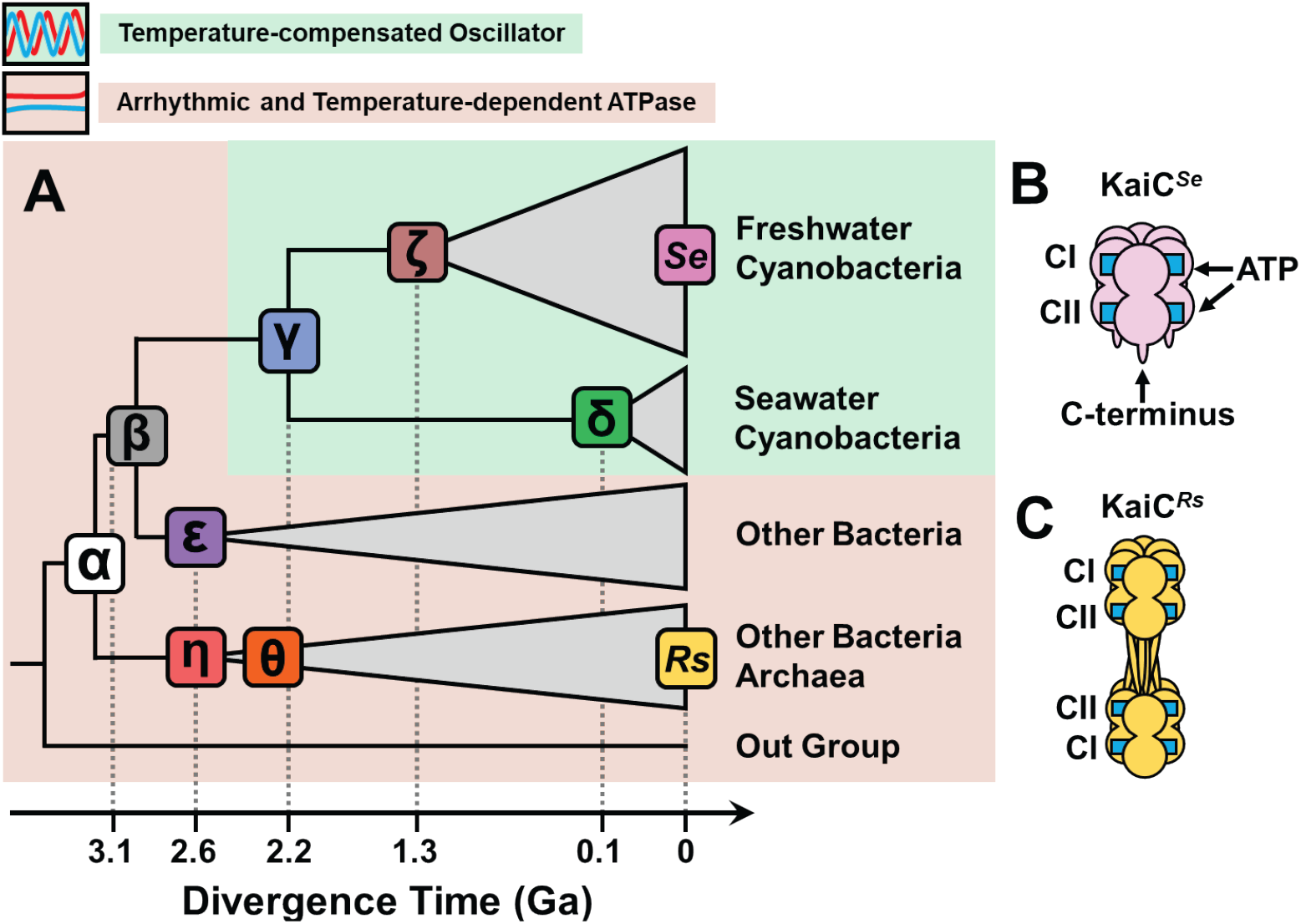
Evolution and diversity of KaiCs. (**A**) A phylogenetic tree constructed using 151 KaiC homologues from extant freshwater cyanobacteria, seawater cyanobacteria, non-cyanobacterial bacteria, and other bacteria including archaea. Seven entries from “α” to “η” represent ancestral KaiCs restored at major nodes between the root and the four clades (11), while “*Se*” and “*Rs*” represent modern KaiCs from *Synechococcus elongatus* and *Rhodobacter sphaeroides*, respectively. The entries displayed on the green background exhibit temperature-compensated and oscillatory activities in the presence of ancestral or species-specific KaiA and KaiB, whereas those placed on the orange background are arrhythmic and temperature-dependent (11, 12). Estimated divergence times for ancestral KaiCs are given in the unit of 1 billion years (= 1 Ga). Schematic drawing of (**B**) KaiC^*Se*^ from *Synechococcus elongatus* PCC7942 (20, 52) and (**C**) KaiC^*Rs*^ from *Rhodobacter sphaeroides* (12).

*Synechococcus elongatus* PCC 7942 (“*Se*” in Fig. 1A) is one of the well-studied freshwater strains (16), and its clock protein KaiC (KaiC^*Se*^) (17) undergoes a temperature-compensated phosphorylation cycle in the presence of KaiA, KaiB, and adenosine triphosphate (ATP) *in vivo* (18) and *in vitro* (19). KaiC^*Se*^ forms a double-ring hexamer upon binding one molecule of ATP to each of an N-terminal (CI) domain and a C-terminal (CII) domain (20) (Fig. 1B). According to the phylogenic tree of a series of existing KaiCs (11), the emergence of the oldest KaiC^α^ dates back approximately 3.1 billion years (Ga) ago (Fig. 1A). KaiC^α^ have evolved through six major branching points (β: 3.1 Ga, γ: 2.2 Ga, δ: 0.1 Ga, ε: 2.6 Ga, ζ: 1.3 Ga, η: 2.6 Ga), and is now diverged into four major clades: freshwater cyanobacteria, marine cyanobacteria, other bacteria, and other bacteria including archaea. A comprehensive functional analysis (11) of these ancestral KaiCs revealed that both the self-sustained rhythmicity and temperature compensation—which were absent in KaiC^α^—emerged during the era of KaiC^γ^ and were subsequently inherited by most modern KaiCs from cyanobacteria in freshwater and seawater (green background in Fig. 1A). Significant molecular evolutions must have occurred in ancient KaiCs to achieve the temperature-compensated cycle, particularly during the transition from KaiC^β^ to KaiC^γ^, but little is known about the structural changes in older KaiCs over approximately 1 Ga since the earliest KaiC^α^ emerged.

Another unresolved issue is the diversity of non-cyanobacterial KaiCs. A major example of such diversity is KaiC from *Rhodobacter sphaeroides* (KaiC^*Rs*^), which evolved though KaiC^η^ (Fig. 1A). A previous study (12) demonstrated that KaiC^*Rs*^ forms a dimer of two hexamers (dodecamer) (Fig. 1C) and functions together with species-specific KaiB (21) as an environmentally responsive hourglass-like timer. However, it remains unclear when and how ancient KaiCs adopted the dodecameric architecture.

In this study, we investigated the shape evolution of ancestral KaiCs using small-angle X-ray scattering (SAXS). The result indicates that the primitive double-ring hexamer was first established in a relatively expanded and asymmetric form and evolved along two contrasting pathways (Fig. 1A). One such pathway is driven by stepwise compactification, leading to the acquisition of the self-sustained rhythmicity and temperature compensation in KaiC^γ^. The other begins with an initial selection of the dodecameric fold at the η point and is subsequently driven by a selection of high ATP consumption and autokinase activity as seen in KaiC^*Rs*^. These observations suggest that the evolutionary history of the Kai-based timing systems cannot be described by a single trajectory, and that the evolution of KaiC holds a potential for further diversification in future.

## Results

### SAXS Measurements of Ancestral and Modern KaiCs

To evaluate the shape evolution, monodispersed KaiC samples were prepared using size-exclusion chromatography (22) and subjected to SAXS measurements at 30°C. Consistent with previous reports (23-25), the scattering intensity *I*(*Q*) of KaiC^*Se*^ decreased monotonously as the angular momentum (*Q*) increased from 0.014 to 0.07 Å^-1^ (Fig. 2A), and exhibited a trough at *Q* of approximately 0.08 Å^-1^ and a peak at *Q* of approximately 0.11 Å^-1^.

**Figure 2.**
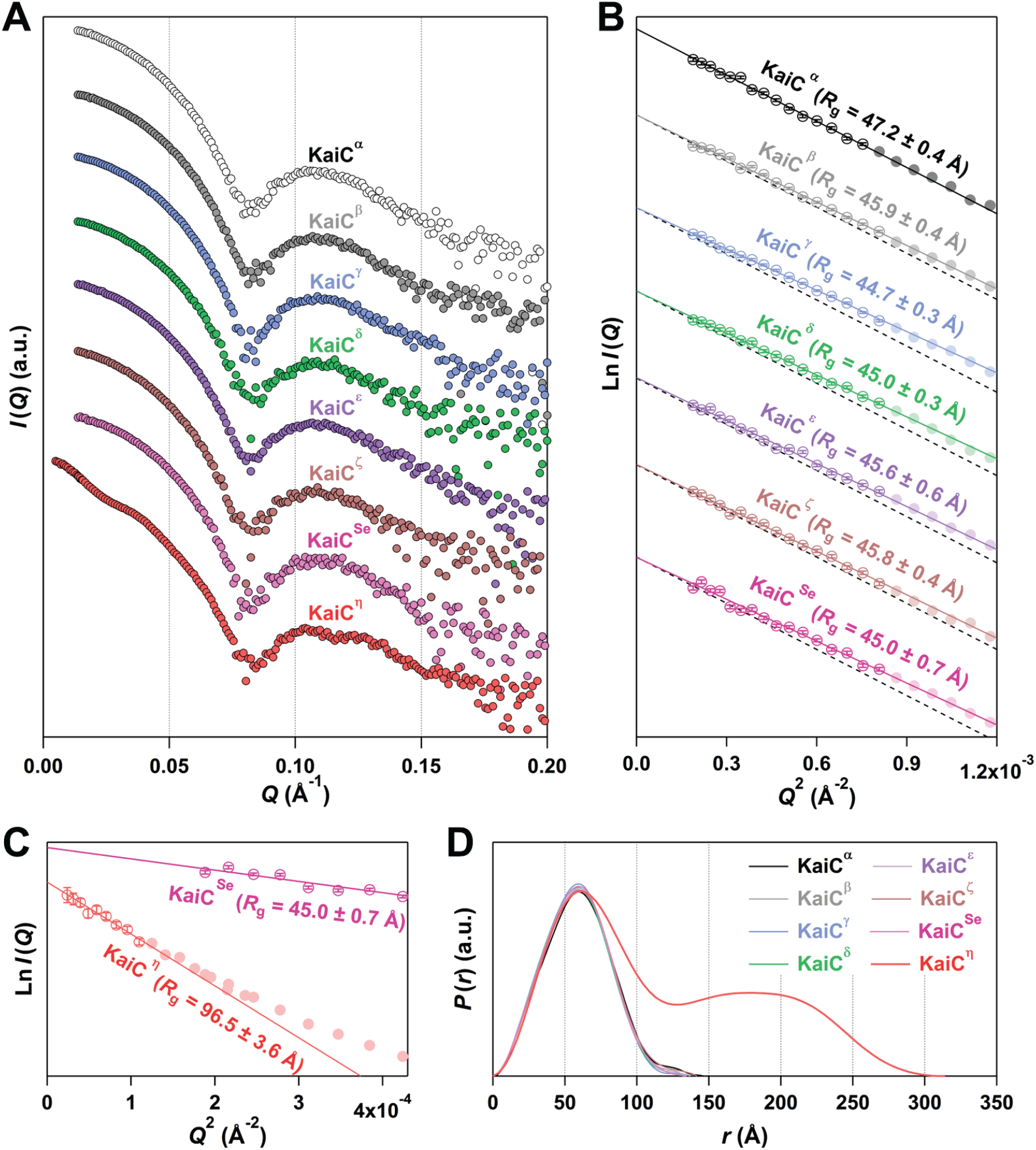
SAXS analyses of ancestral and modern KaiCs at 30°C. (**A**) SAXS intensity, *I*(*Q*), as a function of momentum transfer (*Q*). Each curve is plotted on a logarithmic scale with vertical offsets to ensure clarity of presentation. (**B**) Guinier plots with vertical offsets for clarity of presentation. Each solid line represents linear least-square fit of the experimental data (open circles: *Q*_max_ <1.3 / *R*_g_) to Eq. 1. The *R*_g_ and *I*(0) values determined from its slope and intercept, respectively, are shown in Table 1. Each black dotted line represents a reference line with the slope corresponding to the *R*_g_ value (47.2 Å, black line) of KaiC^α^. (**C**) Guinier plots of KaiC^η^ and KaiC^*Se*^ with vertical offsets for clarity of presentation. Each solid line represents linear least-square fit of the experimental data (open circles: *Q*_max_ <1.0 / *R*_g_ for KaiC^η^ and *Q*_max_ <1.3 / *R*_g_ for KaiC^*Se*^) to Eq. 1. (**D**) Pair distribution function, *P*(*r*). *D*_max_ values determined from *P*(*r*) are summarized in Table 1.

At first glance (Fig. 2A), the SAXS pattern of KaiC^*Se*^ appeared to be shared with most of the ancestral KaiCs: KaiC^α^, KaiC^β^, KaiC^γ^, KaiC^δ^, KaiC^ε^, and KaiC^ζ^ (KaiC^α, β, γ, δ, ε, ζ^). However, these ancestral KaiCs revealed different curvatures of *I*(*Q*) especially in the *Q* range from 0.014 to 0.07 Å^-1^, which were best confirmed by the Guinier analysis shown in Fig. 2B. As demonstrated for oldest KaiC^α^, natural logarithm of *I*(*Q*) in the small *Q* region followed a linear relationship against *Q*^2^, and from its slope (black line in Fig. 2B) the radius of gyration (*R*_g_) of KaiC^α^ was estimated to be 47.2 ± 0.4 Å (Table 1). Second oldest KaiC^β^ showed a smaller *R*_g_ value of 45.9 ± 0.4 Å than KaiC^α^, and KaiC^γ^ located downstream of KaiC^β^ was accompanied by a much smaller *R*_g_ value of 44.7 ± 0.3 Å. These differences from KaiC^α^ can be visually confirmed by the less steeper slopes relative to a reference line (black dotted lines in Fig. 2B) corresponding to the *R*_g_ value of KaiC^α^. While the *R*_g_ values of KaiC^ε^ (45.6 ± 0.6 Å) and KaiC^ζ^ (45.8 ± 0.4 Å) were similar to that of KaiC^β^, the *R*_g_ values of recent KaiC^δ^ (45.0 ± 0.3 Å) and modern KaiC^*Se*^ (45.0 ± 0.7 Å) were as small as that of KaiC^γ^. These observations suggest a tendency for older KaiCs to have larger *R*_g_ values.

**Table 1.**
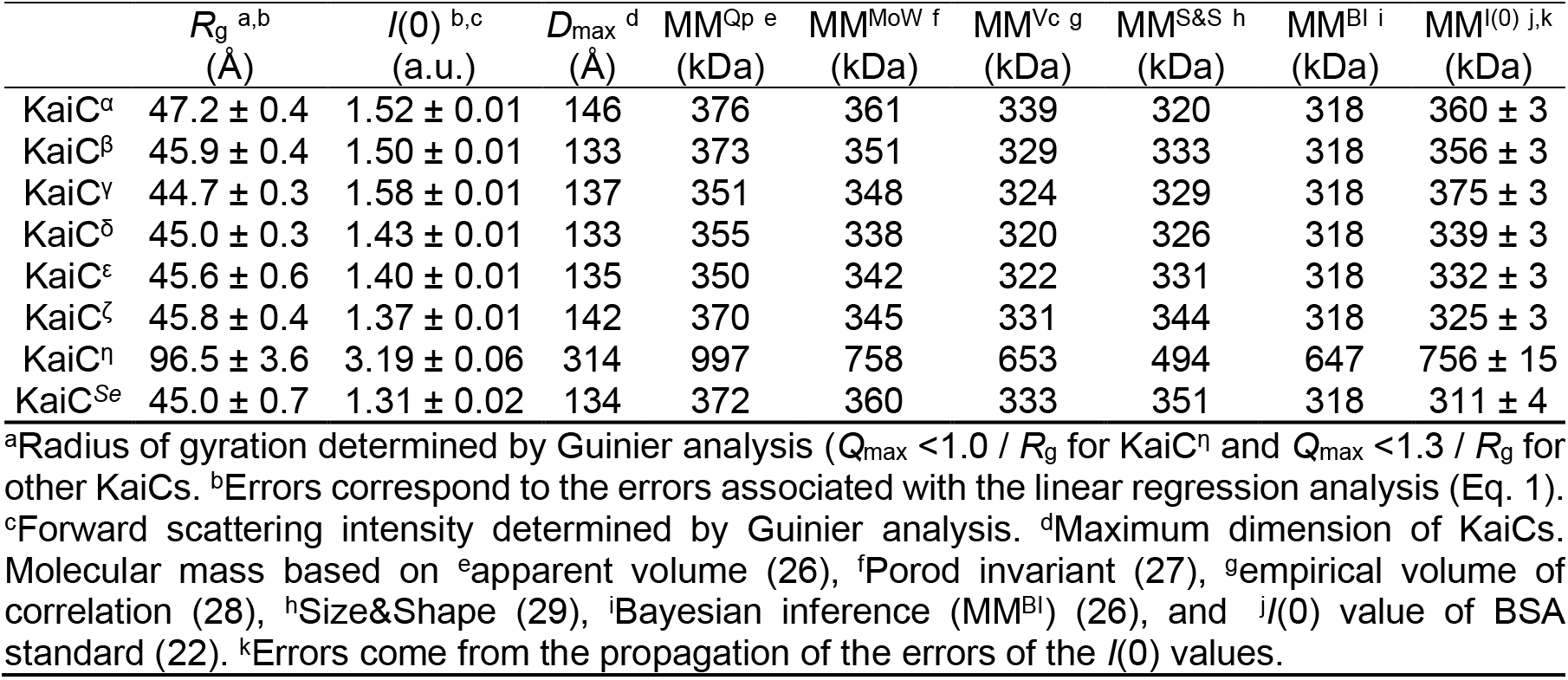
Small-angle X-ray scattering parameters of ancestral and modern KaiCs.

| | $R_g$ <sup>a,b</sup> | $I(0)$ <sup>b,c</sup> | $D_{max}$ <sup>d</sup> | $MM^{Qp}$ <sup>e</sup> | $MM^{MoW}$ <sup>f</sup> | $MM^{Vc}$ <sup>g</sup> | $MM^{S\&S}$ <sup>h</sup> | $MM^{BI}$ <sup>i</sup> | $MM^{I(0)}$ <sup>j,k</sup> |
| --- | --- | --- | --- | --- | --- | --- | --- | --- | --- |
|  | (Å) | (a.u.) | (Å) | (kDa) | (kDa) | (kDa) | (kDa) | (kDa) | (kDa) |
| KaiC<sup>α</sup> | 47.2 ± 0.4 | 1.52 ± 0.01 | 146 | 376 | 361 | 339 | 320 | 318 | 360 ± 3 |
| KaiC<sup>β</sup> | 45.9 ± 0.4 | 1.50 ± 0.01 | 133 | 373 | 351 | 329 | 333 | 318 | 356 ± 3 |
| KaiC<sup>γ</sup> | 44.7 ± 0.3 | 1.58 ± 0.01 | 137 | 351 | 348 | 324 | 329 | 318 | 375 ± 3 |
| KaiC<sup>δ</sup> | 45.0 ± 0.3 | 1.43 ± 0.01 | 133 | 355 | 338 | 320 | 326 | 318 | 339 ± 3 |
| KaiC<sup>ε</sup> | 45.6 ± 0.6 | 1.40 ± 0.01 | 135 | 350 | 342 | 322 | 331 | 318 | 332 ± 3 |
| KaiC<sup>ζ</sup> | 45.8 ± 0.4 | 1.37 ± 0.01 | 142 | 370 | 345 | 331 | 344 | 318 | 325 ± 3 |
| KaiC<sup>η</sup> | 96.5 ± 3.6 | 3.19 ± 0.06 | 314 | 997 | 758 | 653 | 494 | 647 | 756 ± 15 |
| KaiC<sup>Se</sup> | 45.0 ± 0.7 | 1.31 ± 0.02 | 134 | 372 | 360 | 333 | 351 | 318 | 311 ± 4 |

<sup>a</sup>Radius of gyration determined by Guinier analysis ( $Q_{max} < 1.0 / R_g$ for KaiC<sup>η</sup> and $Q_{max} < 1.3 / R_g$ for other KaiCs). <sup>b</sup>Errors correspond to the errors associated with the linear regression analysis (Eq. 1). <sup>c</sup>Forward scattering intensity determined by Guinier analysis. <sup>d</sup>Maximum dimension of KaiCs. <sup>e</sup>Molecular mass based on <sup>e</sup>apparent volume (26), <sup>f</sup>Porod invariant (27), <sup>g</sup>empirical volume of correlation (28), <sup>h</sup>Size&Shape (29), <sup>i</sup>Bayesian inference ( $MM^{BI}$ ) (26), and <sup>j</sup> $I(0)$ value of BSA standard (22). <sup>k</sup>Errors come from the propagation of the errors of the $I(0)$ values.

To examine this tendency in term of oligomerization states, information regarding molecular mass (MM) was extracted from the SAXS data. The *y*-intercept of the linear slope in Fig. 2B corresponds to forward scattering intensity, *I*(0), which is proportional to MM of scatters. As shown in Fig. S1 and Table 1, molecular masses (MM^*I*(0)^) estimated based on the *I*(0) value of bovine serum albumin (BSA) (22) suggest that each of KaiC^α, β, γ, δ, ε, ζ, *Se*^ forms a homo-hexamer (58 kDa × 6 = 348 kDa). The similar results were obtained also from MM estimations using reference-free methods (MM^Qp^ (26), MM^MoW^ (27), MM^Vc^ (28), MM^S&S^ (29), and MM^BI^ (26) in Fig. S1 and Table 1). Thus, the observed differences in the *R*_g_ values among KaiC^α, β, γ, δ, ε, ζ, *Se*^ are attributable to changes in the shape of the hexamers.

The only exception was KaiC^η^ located downstream of KaiC^α^. In contrast to KaiC^α, β, γ, δ, ε, ζ, *Se*^, *I*(*Q*) of KaiC^η^ decreased in a non-monotonous manner in the lower *Q* range and revealed two overlapped peaks at *Q* of approximately 0.105 and 0.125 Å^-1^ (Fig. 2A). The *R*_g_ value of KaiC^η^ was 96.5 ± 3.6 Å (Fig. 2C), which was significantly larger compared to KaiC^α, β, γ, δ, ε, ζ, *Se*^. The MM estimations indicated that KaiC^η^ has a MM of approximately 700 kDa, nearly twice MMs of KaiC^α, β, γ, δ, ε, ζ, *Se*^ (Fig. S1 and Table 1). The unique oligomerization state of KaiC^η^ was suggested also by pair distribution functions, *P*(*r*): a histogram of distances between every pair of two electrons within particles. While each *P*(*r*) of KaiC^α, β, γ, δ, ε, ζ, *Se*^ showed a symmetric peak around 60 Å, that of KaiC^η^ accompanied another peak around 180 Å with a peak area nearly identical to that of the peak around 60 Å (Fig. 2D). Furthermore, the maximum dimension (*D*_max_) more than doubled from 146 to 314 Å upon the divergence of KaiC^α^ into KaiC^η^ (Table 1). These observations suggest that KaiC^η^ is an oligomer consisting of two hexamers separated by roughly 180 Å.

These shape differences among the ancestral and modern KaiCs cannot be simply explained by their phosphorylation status. KaiCs possess dual phosphorylation sites consisting of two adjacent amino acids (S431 and T432 in KaiC^*Se*^) and can exist in four phosphorylation states (30, 31): both are dephosphorylated (DD); only the C-terminal amino acid is phosphorylated (DP); both are phosphorylated (PP); and only the N-terminal amino acid is phosphorylated (PD). Considering that KaiC^*Se*^ becomes less compact upon auto-dephosphorylation (24), the larger *R*_g_ value of KaiC^α^ might be interpreted as resulting from its constitutive accumulation as the DD state (Fig. 3A and *SI Appendix*, Fig. S2). However, despite a similar abundance (approximately 50%) of the fully dephosphorylated DD state in KaiC^γ^ and KaiC^ζ^ (Fig. 3A), they are totally distinct in terms of the compactness (Fig. 3B). The same was observed for another comparison between KaiC^β^ and KaiC^δ^ (Fig. 3A and 3B). Furthermore, the *R*_g_ values were poorly correlated even with the sum of the abundances of the DD and PD states (Fig. 3C), each of which has larger *R*_g_ values in KaiC^*Se*^ than its other states (24). These results suggest that older KaiCs such as KaiC^α, β, ε, ζ, η^ possess distinct and less compact hexamer conformations that cannot be explained solely by their phosphorylation status.

**Figure 3.**
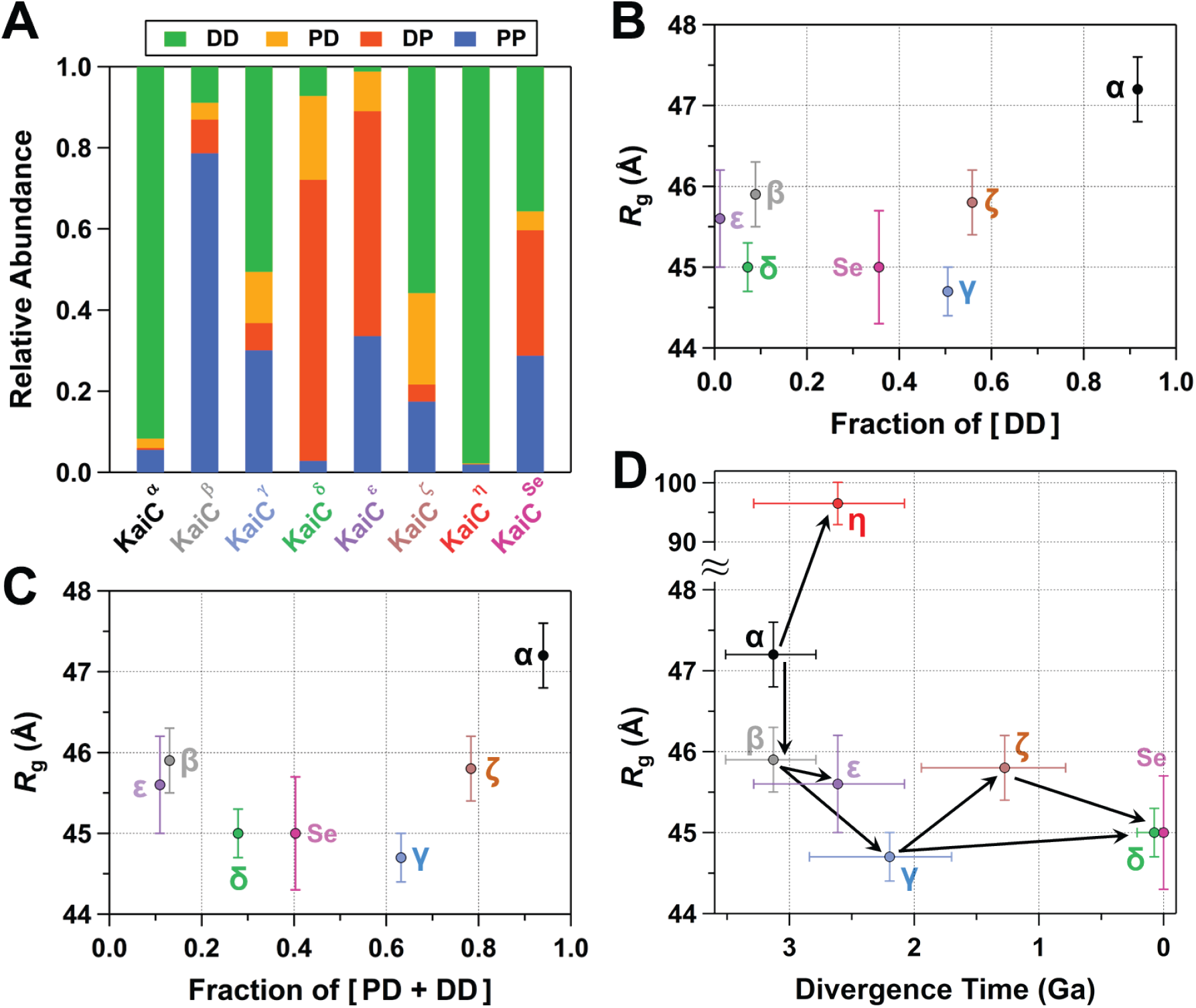
Interpretations of differences in radii of gyration (*R*_g_) among ancestral and modern KaiCs. (**A**) Relative abundances of four phosphorylation states: both of dual phosphorylation sites are dephosphorylated (DD); only the C-terminal amino acid is phosphorylated (DP); both are phosphorylated (PP); and only the N-terminal amino acid is phosphorylated (PD). The fractions were determined by densitometric analysis (51) of gel bands (*SI Appendix*, Fig. S2). Low correlation of *R*_g_ values with the fractions of (**B**) DD state and (**C**) PD + DD states. (**D**) Plot of *R*_g_ values against divergence time of ancestral KaiCs. Black arrows indicate evolutionary pathways of the *R*_g_ values based on Fig. 1A. The divergence times of the ancestral KaiCs were taken from the previous report (11). Errors in the vertical axis originate from the Guinier analysis of the SAXS curves using Eq. 1 (Table 1). Error bars in the horizontal axis correspond to confidence intervals from TimeTree (53) analysis.

A correlation diagram between the *R*_g_ values and divergence times (11) indicates the shape evolution proceeded along two major pathways (Fig. 3D). One is a stepwise compactification pathway, where the *R*_g_ value decreased by approximately 1 Å at each step from KaiC^α^ to KaiC^β^ (approximately 3.1 Ga ago) and then from KaiC^β^ to KaiC^γ^ (approximately 2.2 Ga ago). The other is an oligomerization pathway, where *R*_g_ and *D*_max_ more than doubled in a single step from KaiC^α^ to KaiC^η^ around the time when KaiC^β^ diverged into KaiC^ε^ (approximately 2.6 Ga ago).

### Rigid-body and *Ab Initio* Shape Modellings

To visualize the stepwise compactification pathway, we constructed rigid-body models of KaiC^α, β, γ^ using the crystal structure of KaiC^*Se*^ (32) as a template structure. Independent model building was conducted five times for each KaiC^α, β, γ^ under soft distance constraints for the CI–CI, CII–CII, and CI–CII interfaces (*SI Appendix*, Table S1 and Fig. S3).

In the representative model of KaiC^α^ (Fig. 4A), nearly all the CII domains had substantially reduced contacts with the neighboring CII domains as confirmed by their two-dimensional projections (right, Fig. 4A). Consequently, the overall CII-ring of KaiC^α^ adopted a non-compact and expanded conformation (*SI Appendix*, Fig. S4 and S5). In KaiC^β^, the proportion of the minimally contacting CII domains was slightly reduced, and they coexisted with closely contacting CII domains (Fig. 4B). This co-existence resulted in the partial and asymmetrical expansion of the CII ring of KaiC^β^. In both KaiC^γ^ and KaiC^*Se*^, most of the CII domains were in close contact with the adjacent CII domains, forming nearly symmetrical and compact CII rings (Fig. 4C and 4D). While the theoretical SAXS curve of the rigid-body models (red lines in Fig. 4A–D) agreed with the corresponding experimental SAXS curve (open circles in Fig. 4A–D), each theoretical curve remained clearly distinguishable from the others (upper plots in Fig. 4A–D). The SAXS-based models suggest that KaiC^α^ evolved into KaiC^γ^ via KaiC^β^ with the stepwise compactification of the CII-ring.

**Figure 4.**
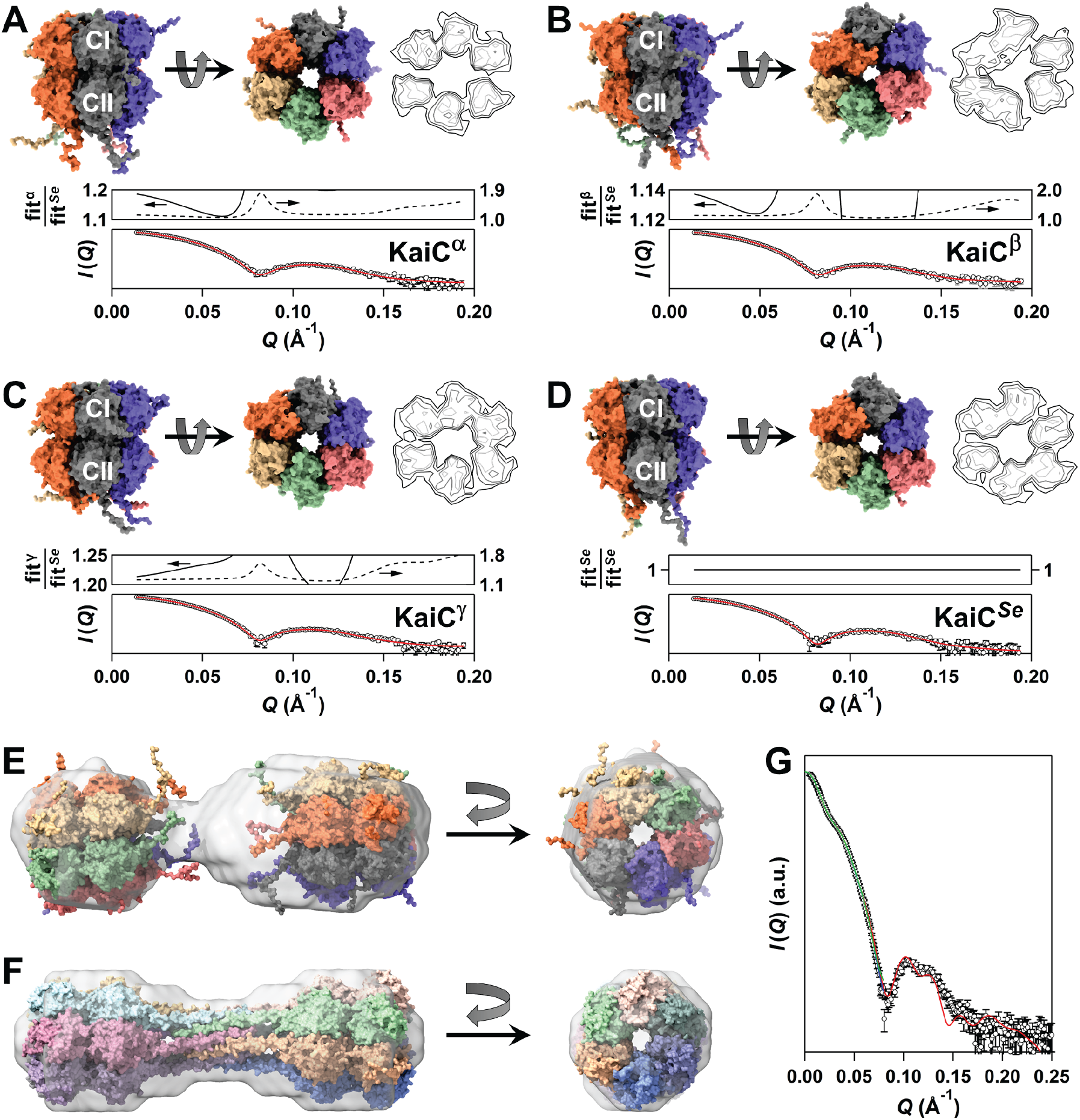
SAXS models for ancestral and modern KaiCs. Rigid-body models of (**A**) KaiC^α^, (**B**) KaiC^β^, (**C**) KaiC^γ^, and (**D**) KaiC^*Se*^ built with CORAL (46). For clarity of presentation, the C-terminal residues (from 501 to 519 or 525) shown in side views (left) are omitted in bottom views (right). Contour images are the vertical projection (voxel spacing of 5 Å) of the CII ring (low: black, medium: gray, high: light gray) (*SI Appendix*, Table S1, Fig. S4, and S5). Open circles and red lines in lower plots correspond to experimental SAXS data and fitting curves of the rigid-body models, respectively. Thick (left axis) and broken lines (right axis) in upper plots indicate fitting curves (fitX: X = α, β, γ, and *Se*) divided by a fitting curve (fitSe) of the KaiC^*Se*^ model. *Ab initio* shapes of KaiC^η^ restored under assumptions of (**E**) P1 and (**F**) P62 symmetries using DAMMIF (47) (*SI Appendix*, Table S1 and Fig. S6). The rigid-body model of KaiC^α^ (panel A) is superimposed on each lobe in the P1 map (panel E). The X-ray crystal structure of KaiC from *Rhodobacter sphaeroides* (KaiC^*Rs*^) (PDB: 8DBA) (12) is superimposed on the P62 map (panel F). (**G**) Experimental SAXS curve of KaiC^η^ (open circles), and theoretical SAXS curves of the P1 envelope shown in panel E (blue line, *χ*^2^ = 1.09 ± 0.04), the P62 envelope shown in panel F (green line, *χ*^2^ = 2.45 ± 0.62), and the X-ray crystal structure of KaiC^*Rs*^ shown in panel F (red line, *χ*^2^ = 4.657 with CRYSOL (54)).

To visualize the oligomerization pathway, *ab initio* shapes of KaiC^η^ were restored from its experimental SAXS curve (Fig. 2A) under the assumption of P1 or P62 symmetry (*SI Appendix*, Fig. S6). The P1 reconstruction yielded a bilobed envelope with the longest dimension of 315 Å (Fig. 4E). While the size of two lobes was asymmetrical along with the major axis direction (left, Fig. 4E), each lobe possessed a volume enough to accommodate one hexameric rigid-body model of KaiC^α^ (right, Fig. 4E). However, because the centers of the two lobes are separated by approximately 180 Å, the two KaiC^α^ hexamers placed on the map are not sufficiently close to interact with each other.

On the other hand, the P62 reconstruction generated an envelope with sufficient density in the linker region connecting the two lobes (Fig. 4F). The envelope showed a good overlap with the X-ray crystal structure of KaiC^*Rs*^: the modern KaiC homolog located downstream of KaiC^η^ (Fig. 1A and 1C). KaiC^*Rs*^ is known to homodimerize through a coiled-coil domain in its C-terminal region that is approximately 50 amino acids longer than KaiC^α^ and KaiC^*Se*^ (12). Because the C-terminus of KaiC^η^ is as long as that of KaiC^*Rs*^ (*SI Appendix*, Fig. S7), it is reasonable to conclude that the two hexamers of KaiC^η^ dimerize in a manner similar to KaiC^*Rs*^ through interactions between their C-termini. To be consistent with these observations, the experimental SAXS curve of KaiC^η^ could be fitted reasonably with the theoretical SAXS curve of KaiC^*Rs*^ (Fig. 4G). Another shape-evolution pathway followed by earliest KaiC^α^ is likely driven by a selective pressure favoring the tail-to-tail dimerization of the two hexamers.

### Evolution of Active Sites as Dumbbell-shaped Dodecameric KaiCs

Although KaiC^η^ and KaiC^*Rs*^ share the dumbbell-shaped structure (Fig. 4F), there is a more than 50-fold difference in their ATP consumption rates. At 30°C, KaiC^η^ consumes 4 ATP per day (11), which is the sum of the ATPase activity (33, 34) at the CI site (CI-ATPase) and the combined ATPase (35), kinase (34), and phosphatase (36) activities at the CII site. In KaiC^*Rs*^, the CI and CII active sites exhibit roughly equivalent activity, and their total activity results in a higher consumption rate of approximately 220 ATP d^-1^ compared to other KaiC homologs (12). During the approximately 2.6 Ga evolutionary process from KaiC^η^ to KaiC^*Rs*^, therefore, the activities of both the CI and CII sites increased by at least 28-fold.

To identify the timing when the activity increased and factors behind it, we revisited the evolutionary linage from KaiC^η^ to KaiC^*Rs*^ using the previously reported phylogenic tree (*SI Appendix*, Fig. S8) (11). We found that several key residues involved in the active sites had changed from the KaiC^η^-type to the KaiC^*Rs*^-type (*SI Appendix*, Fig. S7) at a divergence point (θ) corresponding to 2.3 (2.9–1.8: upper–lower confidence intervals) Ga ago (Fig. 1A and *SI Appendix*, Fig. S8). To visualize the evolution of the active sites during the oligomerization pathway, we constructed ADP·Mg^2+^-bound models of KaiC^η^ and KaiC^θ^ using AlphaFold3 (37), and compared them with the X-ray crystal structure of KaiC^*Rs*^ (PDB: 8DBA).

The resulting models predicted that the CI active site had loosened at the θ node. In the predicted model of KaiC^η^, two adjacent CI domains were closely interacted through a hydrogen-bonding network formed by E175, S178, and S179 (Fig. 5A). It was predicted that this hydrogen-bonding network was first weakened in KaiC^θ^ due to the loss of two consecutive serine residues at 178th and 179th positions in KaiC^η^ (Fig. 5B), and was subsequently disrupted in KaiC^*Rs*^ by the substitution of E179 (KaiC^θ^) with D171 (KaiC^*Rs*^) having the shorter side-chain length (Fig. 5C). According to previous studies on KaiC^*Se*^, the CI-ATPase cycle consists of multiple processes, including the positioning of a water molecule attacking ATP, nucleotide exchanges, and the release of an inorganic phosphate; each of these processes is closely coupled with structural changes at the CI–CI interface (33, 35). Thus, the loosening of the CI–CI interface during the transition from KaiC^η^ to KaiC^θ^ and then to KaiC^*Rs*^ may have contributed the efficient exchange of CI-ADP and release of the inorganic phosphate.

**Figure 5.**
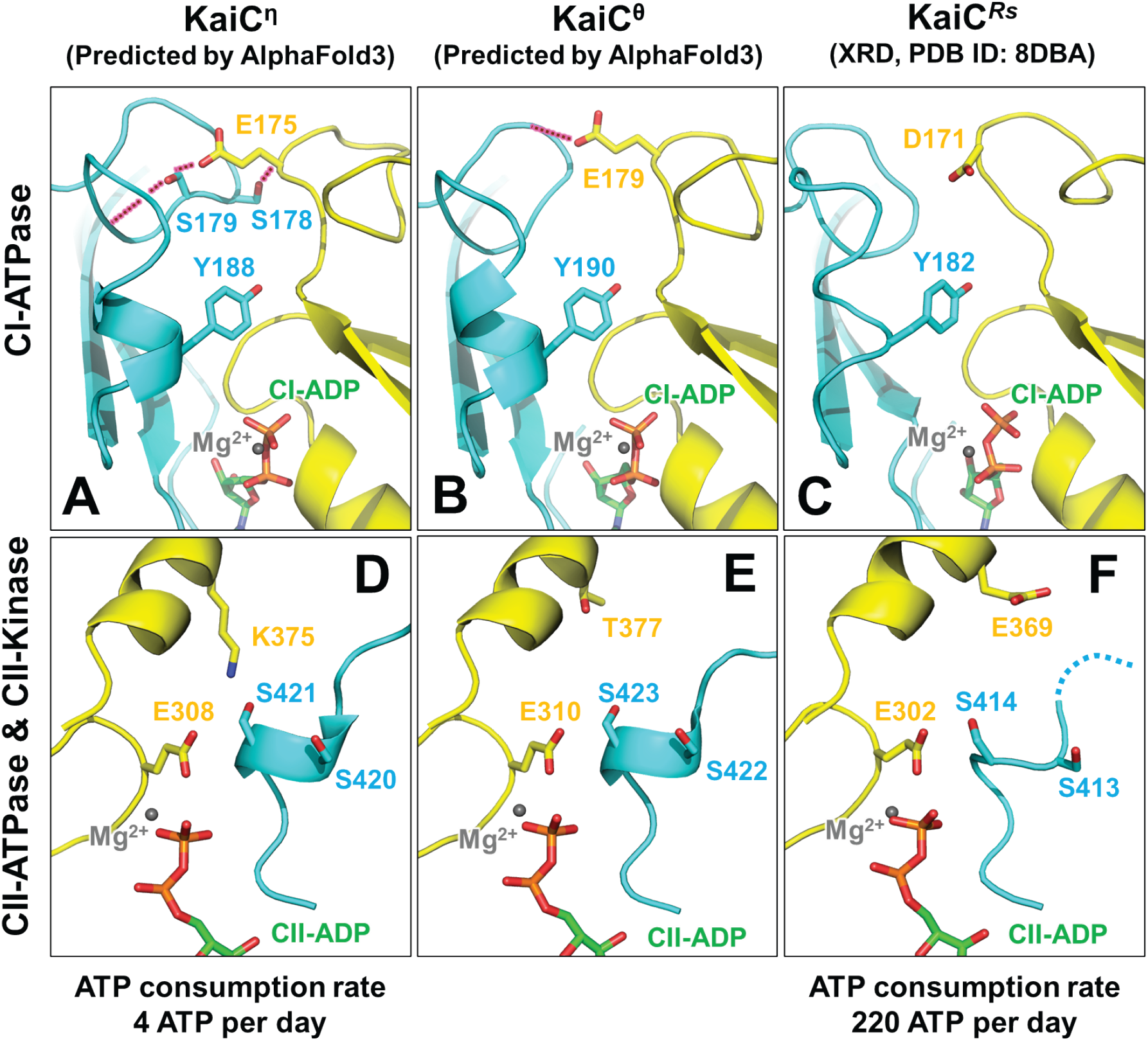
Comparison of predicted structures of KaiC^η^ and KaiC^θ^ using AlphaFold3 with the X-ray crystal structures of KaiC^*Rs*^ (12). Active sites in the CI domains of (**A**) KaiC^η^, (**B**) KaiC^θ^, and (**C**) KaiC^*Rs*^. Red dotted lines represent potential hydrogen-bonding networks. Active sites in the CII domains of (**D**) KaiC^η^, (**E**) KaiC^θ^, and (**F**) KaiC^*Rs*^. Catalytic glutamates: E308 in KaiC^η^, E310 in KaiC^θ^, and E302 in KaiC^*Rs*^. Dual phosphorylation sites: S420 and S421 in KaiC^η^, S422 and S423 in KaiC^θ^, and S413 and S414 in KaiC^*Rs*^.

A notable structural difference was predicted in the CII active site, particularly in a residue that could influence the CII-ATP hydrolysis and autophosphorylation. In KaiC^η^, K375 was located near a catalytic Glu (E308) and a phosphorylation site (S421), close enough to affect their electrostatic properties (Fig. 5D). The amino acid corresponding to K375 in KaiC^η^ is R385 in KaiC^*Se*^ (*SI Appendix*, Fig. S7). In KaiC^*Se*^, R385 is known to inhibit CII-ATPase and CII-kinase by neutralizing a catalytic Glu (E318 in KaiC^*Se*^) and inactivating its ability to subtract protons from a lytic water or phosphorylation sites (38). Therefore, it is reasonable to suggest that the catalytic activity of E308 is downregulated by K375 in KaiC^η^. It was predicted the catalytic Glu was released from the inhibitory neutralization in KaiC^θ^ by the substitution of K375 (KaiC^η^) with T377 (KaiC^θ^) (Fig. 5E), and was subsequently placed under a negative electrostatic environment in KaiC^*Rs*^ by the replacement of T377 (KaiC^θ^) with E369 (KaiC^*Rs*^) (Fig. 5F). To be consistent with this interpretation, KaiC^η^ and KaiC^*Rs*^ are constitutively dephosphorylated (Fig. 3A) and phosphorylated (12), respectively.

The predicted structures suggest that KaiC^η^ underwent the local structural changes in both the CI and CII active sites within the dodecameric shape, resulting in the formation of the highly active KaiC^*Rs*^.

## Discussion

The shape evolution of KaiC^α^, the oldest double-ring predecessor, proceeded along the two contrasting pathways (Fig. 6). One is the pathway involving the stepwise compactification that was a necessary event for ancient KaiCs to evolve into the self-sustained and temperature-compensated oscillator. Since both KaiC^α^ and KaiC^β^ are arhythmic in the presences of ancestral KaiAs and KaiBs (11) (Fig. 1A), the initial reduction by approximately 1 Å in *R*_g_ can be considered insufficient to confer the autonomous rhythmicity to KaiC^β^. This is also supported by the crystal structure of KaiC^β^, in which the CII ring is crystallized in an asymmetrical and less compact conformation (*SI Appendix*, Fig. S9A and S9B) (11). The CII ring of KaiC^β^ in the crystal (*SI Appendix*, Fig. S9A) is less asymmetric than in solution phase (Fig. 4B and *SI Appendix*, Fig. S4B), but this will be an inevitable consequence of confining the molecules in the crystal lattice. Conversely, the asymmetry of KaiC^β^ can be considered yet pronounced in solution that it does not disappear completely even under confinement by the crystal lattice. This insufficiently compacted shape of KaiC^β^ was inherited by not self-sustained KaiC^ε^ (Fig. 6) and eventually passed on to the downstream non-cyanobacterial lineage (Fig. 1A).

**Figure 6.**
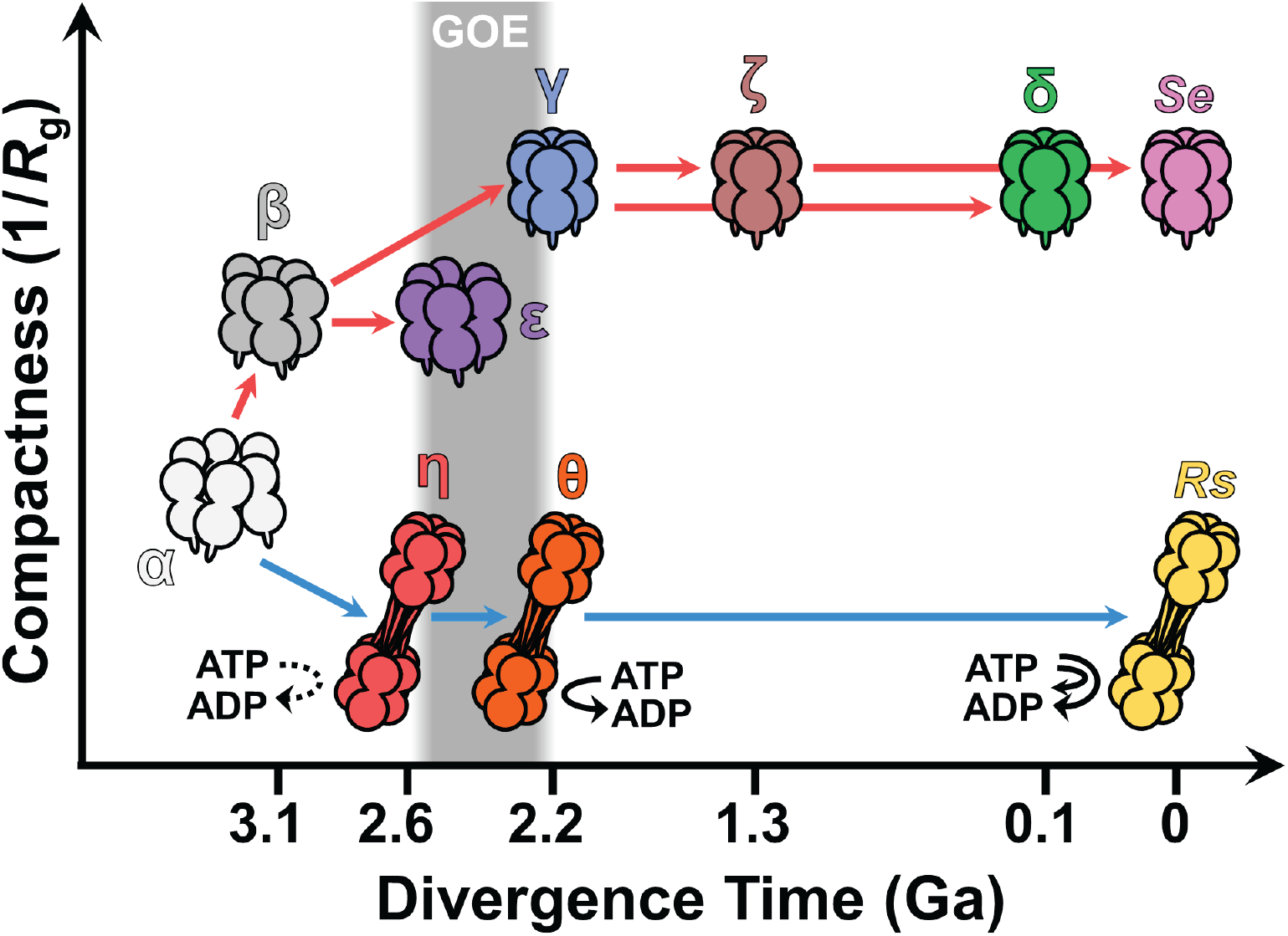
Shape evolution of KaiCs along stepwise compactification (red arrows) and oligomerization (blue arrows) pathways. GOE: Great Oxidation event (40).

KaiC^γ^ emerged as the clock component of the self-sustained and temperature-compensated oscillator after the further compactification from KaiC^β^ to KaiC^γ^ (Fig. 6), that is, after the net decrease in *R*_g_ by approximately 2 Å compared to KaiC^α^ (Fig. 3D). The more compact nature of KaiC^γ^ than KaiC^β^ is also supported by their crystal structures. The CII ring in the crystal structure of KaiC^γ^ exhibits a symmetrical and compact conformation, similar to KaiC^*Se*^ and KaiC^δ^ (*SI Appendix*, Fig. S9C and S9D) (11). On the other hand, in KaiC^*Se*^, point mutations that only impair temperature compensation while preserving the rhythmicity have been identified (39). Therefore, it is not necessarily clear to what extent the compactification in KaiC^γ^ contributed its acquisition of temperature compensation. It cannot be ruled out that the temperature insensitivity of KaiC^γ^ is a byproduct of becoming the component of the autonomous oscillator. In any case, modern KaiCs from freshwater cyanobacteria, including KaiC^*Se*^, inherited the compact, self-sustained, and temperature-compensated nature of KaiC^γ^ via KaiC^ζ^, and in parallel, extant KaiCs from marine cyanobacteria inherited the similar properties via KaiC^δ^ (Fig. 6).

The other is the pathway that leads to a loss of compactness. Kern and colleagues (12) discovered the dimer of two extant KaiC^*Rs*^ hexamers and postulated that the dimerization could have occurred no later than 1 Ga ago based on their phylogenic analysis. The present SAXS study on KaiC^η^ provides the direct experimental evidence supporting their prediction.

KaiC^η^ emerged as the dumbbell-shaped dodecamer 2.6 Ga ago (Fig. 6), but its ATP consumption rate (11) and autokinase activity (Fig. 3A) remained as low as KaiC^α^. According to the results of predicting the structural evolution from KaiC^η^ to KaiC^*Rs*^ (Fig. 5), the evolution of the CI and CII active sites continued for a certain period after the formation of the dodecameric fold, leading to the establishment of the KaiC^*Rs*^-like active sites in KaiC^θ^ approximately 2.3 Ga ago (Fig. 6 and *SI Appendix*, Fig. S8). It appears that the oligomerization pathway was driven by the sequential action of two selective pressures: one favoring the dimerization between the C-terminal tails of the two hexamers, and the other favoring the higher ATP consumption and autokinase activity (Fig. 6).

We believe that it is the latter selective pressure that has granted KaiC^*Rs*^ its high environmental adaptability. KaiC^*Rs*^, together with species-specific KaiB (21), functions as an hourglass-like clock system whose phosphorylation state is reset in response to environmental signals such as the ATP-to-ADP ratio (12). The higher ATP consumption and kinase activity should shorten the time lag between the reception of day-night signals and the system’s response, thereby contributing to the establishment of a more responsive passive timing system.

In summary, the present study proposes a plausible scenario for the shape evolution of KaiCs on Earth from ancient and modern times. In approximately 1 Ga since KaiC^α^ first appeared, its shape had undergone dramatic evolution and given rise to the two distinct prototypes of extant KaiCs (Fig. 6). The timing of this drastic shape evolution coincides the Great Oxidation event (GOE) (Fig. 6), when the concentration of free oxygen first rose on Earth (40). Thus, the modern time-keeping systems are products of the adaptive shape evolution to geological fluctuations at the era of GOE. Our study clearly demonstrates that the stepwise compactification pathway is one that gave rise to the self-sustained and temperature-compensated Kai oscillators.

## Materials and Methods

### Preparation of Modern and Ancestral KaiCs

According to the previous study (11, 36, 41, 42), modern KaiC^*Se*^ and ancestral KaiC^α^, KaiC^β^, KaiC^γ^, KaiC^δ^, KaiC^ε^, KaiC^ζ^, and KaiC^η^ were expressed in *E. coli* BL21 cells and then purified by affinity chromatography using either glutathione S-transferase tag or histidine tag. The eluted samples were further purified using size exclusion chromatography and ion exchange chromatography.

### SAXS Experiments

SAXS experiments were conducted using Nano-Viewer (RIGAKU) equipped with a microfocus rotating anode X-ray generator, RA-Micro7 (Rigaku). Scattering images were recorded with a hybrid pixel array detector, PILATUS 200K (DECTRIS).

Immediately before the SAXS measurements, purified KaiC samples were passed through buffer-equilibrated gel filtration column (Superdex 200 Increase 10/300, Cytiva) to remove aggregates if any. For modern KaiCs including KaiC^*Se*^, we used a buffer containing 50 mM Tris-HCl (pH8), 150 mM NaCl, 5 mM ATP, 5 mM MgCl_2_, 0.5 mM ethylenediaminetetraacetic acid (EDTA), and 3 mM dithiothreitol (DTT). For ancestral KaiCs other than KaiC^*Se*^, we used a buffer containing 50 mM Tris-HCl (pH7), 150 mM NaCl, 5 mM ATP, 5 mM MgCl_2_, 0.5 mM EDTA, and 3 mM DTT. The cuvette was filled with the mono-dispersed fraction after the gel filtration and placed on a Peltier temperature-controlling cuvette holder maintained at 30 ± 0.1°C.

Depending on the protein concentration (1.5∼6.0 mg/ml), 5 to 20 consecutive frames of 6 min exposure to the X-ray wavelength (*λ*) of 1.5418 Å were recorded at a detector distance of 715 mm. Only radiation–damage-free frames were circularly averaged and normalized by the total irradiation time and the protein concentration (*c*: w/v) to obtain a scattering intensity, *I*(*Q, c*), where *Q* is defined by 4πsin(*θ*) / *λ* and 2*θ* is the scattering angle. For KaiC^η^, 10 successive frames of 10 min exposure were recorded at another detector distance of 1,004 mm to cover the small *Q* region from 0.004888 to 0.009776 Å^-1^.

### SAXS Data Analysis

Experimental SAXS data were processed using a program suite, ATSUS 4.0 (43). The dataset of *I*(*Q, c*) measured at different protein concentrations were extrapolated to infinite dilution to estimate *I*(*Q*) using a software, PRIMUS (44). Solely in the case of KaiC^η^, *I*(*Q, c*) collected at the detector distance of 1,004 mm were nearly unchanged in the protein concentrations from 2.6 to 3.8 mg/ml and thus averaged as *I*(*Q*) without extrapolating to infinite dilution. The resultant *I*(*Q*) of KaiC^η^ was then merged with the *I*(*Q*) collected at the detector distance of 715 mm using a software, datmerge (43). Pair distribution functions, *P*(*r*), were calculated from *I*(*Q*) using a software, GNOM (45).

The forward-scattering intensity, *I*(0), and the radius of gyration, *R*_g_, were determined by the Guinier analysis of *I*(*Q*) using the following equation.

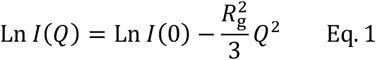

Since *I*(0) is approximately proportional to MM of a protein, MMs of KaiCs were determined using the *I*(0) value of bovine serum albumin (BSA, Sigma #A7638) as a standard (22). At the same time, molecular masses were estimated using concentration-independent methods based on the Porod invariant (MM^MoW^) (27), apparent volume obtained from the Porod invariant (MM^Qp^) (26), empirical volume of correlation (MM^Vc^) (28), Size&Shape (MM^S&S^) (29), and Bayesian inference (MM^BI^) (26).

### SAXS-based Modeling

Rigid-body models of KaiC^α^, KaiC^β^, KaiC^γ^, and KaiC^*Se*^ were built using CORAL (46). The crystal structure of KaiC^*Se*^ (PDB: 2GBL) (32) was used as a template model to construct initial structures, in which each protomer was treated as a flexible N-terminus of thirteen residues, a CI domain of residues 14–246, a flexible linker of nine residues, a CII domain of residues 255–500, and a flexible linker of residues 501–519 (for KaiC^*Se*^ and KaiC^γ^) or 501–525 (6xhis-tagged KaiC^α^ and KaiC^β^) (*SI Appendix*, Table S1). To explore the conformational space of hexamers effectively, we imposed soft distance constraints (a penalty value of 10) on three pairs of C_α_ atoms: 9.5 Å for the C_α_ distance between F199 and S146 in the CI–CI interface, 7.3 Å for T432 and A382 in the CII–CII interface, and 6.6 Å for P236 and E357 in the CI and CII domains, respectively (*SI Appendix*, Fig. S3). Each of these three distances correspond to the corresponding distance in the crystal structure of KaiC^*Se*^ (PDB: 2GBL) (32). Each model shown in Fig. 4 is the representative model from five independent calculations (*SI Appendix*, Table S1 and Fig. S4).

*Ab initio* shapes of KaiC^η^ were reconstructed using DAMMIF (47) under the assumption of P1 or P62 symmetry. Twenty independent reconstructions were performed for each symmetry (*SI Appendix*, Table S1). The reconstruction under P1 symmetry resulted in three classes: a major class (12/20), two minor classes (6/20 and 2/20) similar to the major class (*SI Appendix*, Fig. S6A). The reconstruction under P62 symmetry revealed three classes: a major class (13/20), two minor classes (4/20 and 3/20) distinct from the major class (*SI Appendix*, Fig. S6B). The reconstructed models in the major class were averaged using DAMAVER (48) and visualized in Fig. 4 with Situs package (49) and ChimeraX (50).

### KaiC Phosphorylation Assay

An aliquot of every KaiC sample prepared for SAXS experiments was mixed with Laemmli sample buffer and then subjected to sodium dodecyl sulfate polyacrylamide gel electrophoresis analysis (30). Fractions of the phosphorylated species of KaiC were determined by densitometric analysis of gel bands (51).

### Structural Prediction using AlphaFold3

The amino acid sequences of KaiC^η^ and KaiC^θ^ (11) were submitted to AlphaFold server (37). Because their dodecameric assembly exceeded the maximum size allowed, hexameric models were predicted instead. This simplification is reasonable, as the structure of the hexamer of KaiC^*Rs*^ without its C-terminal tail (PDB: 8DB3) is essentially identical to the hexamers constituting the dodecamer (PDB: 8DBA). The active sites of KaiC^*Rs*^ were reconstructed by incorporating twelve molecules of ADP and twelve Mg^2+^ ions. Five candidate models were generated for each of KaiC^η^ and KaiC^θ^. The five models were essentially similar with variations limited to flexible loops and side chains. A representative model with the highest confidence score was selected for presentation in Fig. 5.

## Acknowledgments

This study was supported by JSPS Grants-in-Aid for Scientific Research (22H04984 to S.A., 24H02301 to S.A., and 23H02448 to S.A., and 26H01830 to Y.F.) and partly by Takeda Science Foundation (to S.A.) and Toyoaki Scholarship Foundation (to S.A.).

## References

1. M. J. Harms, J. W. Thornton, Evolutionary biochemistry: revealing the historical and physical causes of protein properties. Nat. Rev. Genet. 14, 559–571 (2013).

2. Y. Raynes, P. D. Sniegowski, Experimental evolution and the dynamics of genomic mutation rate modifiers. Heredity 113, 375–380 (2014).

3. Q. Y. Tang, W. T. Ren, J. Wang, K. Kaneko, The Statistical Trends of Protein Evolution: A Lesson from AlphaFold Database. Mol. Biol. Evol. 39, msac197 (2022).

4. M. L. Jabbur, C. H. Johnson, Spectres of Clock Evolution: Past, Present, and Yet to Come. Front. Physiol. 12, 815847 (2022).

5. R. S. Edgar et al., Peroxiredoxins are conserved markers of circadian rhythms. Nature 485, 459–464 (2012).

6. V. Dvornyk, O. Vinogradova, E. Nevo, Origin and evolution of circadian clock genes in prokaryotes. Proc. Natl. Acad. Sci. U.S.A. 100, 2495–2500 (2003).

7. M. M. Liao et al., The P-loop NTPase RUVBL2 is a conserved clock component across eukaryotes. Nature 642, 165–173 (2025).

8. N. Kon et al., Na+/Ca2+ exchanger mediates cold Ca2+ signaling conserved for temperature-compensated circadian rhythms. Sci. Adv. 7, eabe8132 (2021).

9. M. A. Woelfle, O. Y. Yan, K. Phanvijhitsiri, C. H. Johnson, The adaptive value of circadian clocks: An experimental assessment in cyanobacteria. Curr. Biol. 14, 1481–1486 (2004).

10. S. L. Li et al., Reconstruction of the ancient cyanobacterial proto-circadian clock system KaiABC. EMBO J. 44, 3025–3046 (2025).

11. A. Mukaiyama et al., Evolutionary origins of self-sustained Kai protein circadian oscillators in cyanobacteria. Nat. Commun. 16, 4541 (2025).

12. W. Pitsawong et al., From primordial clocks to circadian oscillators. Nature 616, 183–189 (2023).

13. N. M. Schmelling et al., Minimal tool set for a prokaryotic circadian clock. BMC Evol. Biol. 17, 169 (2017).

14. C. S. Pittendrigh, Temporal Organization - Reflections of a Darwinian Clock-Watcher. Annu. Rev. Physiol. 55, 16–54 (1993).

15. C. Köbler et al., Two KaiABC systems control circadian oscillations in one cyanobacterium. Nat. Commun. 15, 7674 (2024).

16. M. X. Fang, C. L. Partch, A. Liwang, S. S. Golden, Prokaryotic Circadian Systems: Cyanobacteria and Beyond. Annu. Rev. Microbiol. 79, 523–545 (2025).

17. M. Ishiura et al., Expression of a gene cluster as a circadian feedback process in cyanobacteria. Science 281, 1519–1523 (1998).

18. J. Tomita, M. Nakajima, T. Kondo, H. Iwasaki, No transcription-translation feedback in circadian rhythm of KaiC phosphorylation. Science 307, 251–254 (2005).

19. M. Nakajima et al., Reconstitution of circadian oscillation of cyanobacterial KaiC phosphorylation in vitro. Science 308, 414–415 (2005).

20. R. Pattanayek et al., Visualizing a circadian clock protein: Crystal structure of KaiC and functional insights. Mol. Cell. 15, 375–388 (2004).

21. H. K. Wayment-Steele et al., The conformational landscape of fold-switcher KaiB is tuned to the circadian rhythm timescale. Proc. Natl. Acad. Sci. U.S.A. 121, e2412293121 (2024).

22. S. Akiyama, Quality control of protein standards for molecular mass determinations by small-angle X-ray scattering. J. Appl. Crystallogr. 43, 237–243 (2010).

23. S. Akiyama, A. Nohara, K. Ito, Y. Maéda, Assembly and disassembly dynamics of the cyanobacterial periodosome. Mol. Cell. 29, 703–716 (2008).

24. Y. Murayama et al., Tracking and visualizing the circadian ticking of the cyanobacterial clock protein KaiC in solution. EMBO J. 30, 68–78 (2011).

25. Y. Yunoki et al., Overall structure of fully assembled cyanobacterial KaiABC circadian clock complex by an integrated experimental-computational approach. Commun. Biol. 5, 184 (2022).

26. N. R. Hajizadeh, D. Franke, C. M. Jeffries, D. I. Svergun, Consensus Bayesian assessment of protein molecular mass from solution X-ray scattering data. Sci. Rep. 8, 7204 (2018).

27. H. Fischer, M. D. Neto, H. B. Napolitano, I. Polikarpov, A. F. Craievich, Determination of the molecular weight of proteins in solution from a single small-angle X-ray scattering measurement on a relative scale. J. Appl. Crystallogr. 43, 101–109 (2010).

28. R. P. Rambo, J. A. Tainer, Accurate assessment of mass, models and resolution by small-angle scattering. Nature 496, 477–481 (2013).

29. D. Franke, C. M. Jeffries, D. I. Svergun, Machine Learning Methods for X-Ray Scattering Data Analysis from Biomacromolecular Solutions. Biophys. J. 114, 2485–2492 (2018).

30. T. Nishiwaki et al., A sequential program of dual phosphorylation of KaiC as a basis for circadian rhythm in cyanobacteria. EMBO J. 26, 4029–4037 (2007).

31. M. J. Rust, J. S. Markson, W. S. Lane, D. S. Fisher, E. K. O’Shea, Ordered phosphorylation governs oscillation of a three-protein circadian clock. Science 318, 809–812 (2007).

32. R. Pattanayek et al., Analysis of KaiA-KaiC protein interactions in the cyano-bacterial circadian clock using hybrid structural methods. EMBO J. 25, 2017–2028 (2006).

33. J. Abe et al., Atomic-scale origins of slowness in the cyanobacterial circadian clock. Science 349, 312–316 (2015).

34. K. Terauchi et al., ATPase activity of KaiC determines the basic timing for circadian clock of cyanobacteria. Proc. Natl. Acad. Sci. U.S.A. 104, 16377–16381 (2007).

35. Y. Furuike et al., Regulation mechanisms of the dual ATPase in KaiC. Proc. Natl. Acad. Sci. U.S.A. 119, e2119627119 (2022).

36. T. Nishiwaki, T. Kondo, Circadian Autodephosphorylation of Cyanobacterial Clock Protein KaiC Occurs via Formation of ATP as Intermediate. J. Biol. Chem. 287, 18030–18035 (2012).

37. J. Abramson et al., Accurate structure prediction of biomolecular interactions with AlphaFold 3. Nature 630, 493–500 (2024).

38. Y. Furuike, Y. Onoue, S. Saito, T. Mori, S. Akiyama, The priming phosphorylation of KaiC is activated by the release of its autokinase autoinhibition. PNAS Nexus 4, pgaf136 (2025).

39. Y. Furuike et al., Cross-scale analysis of temperature compensation in the cyanobacterial circadian clock system. Commun. Phys. 5, 75 (2022).

40. A. Bekker et al., Dating the rise of atmospheric oxygen. Nature 427, 117–120 (2004).

41. A. Mukaiyama, D. Y. Ouyang, Y. Furuike, S. Akiyama, KaiC from a cyanobacterium sp. PCC 7428 retains functional and structural properties required as the core of circadian clock system. Int. J. Biol. Macromol. 131, 67–73 (2019).

42. D. Y. Ouyang et al., Development and Optimization of Expression, Purification, and ATPase Assay of KaiC for Medium-Throughput Screening of Circadian Clock Mutants in Cyanobacteria. Int. J. Mol. Sci. 20, 2789 (2019).

43. D. Franke, T. Gräwert, D. I. Svergun, New features in ATSAS-4, a program suite for smallangle scattering data analysis. J. Appl. Crystallogr. 58, 1027–1033 (2025).

44. K. Manalastas-Cantos et al., ATSAS 3.0: expanded functionality and new tools for small-angle scattering data analysis. J. Appl. Crystallogr. 54, 343–355 (2021).

45. D. I. Svergun, Determination of the Regularization Parameter in Indirect-Transform Methods Using Perceptual Criteria. J. Appl. Crystallogr. 25, 495–503 (1992).

46. M. V. Petoukhov et al., New developments in the program package for small-angle scattering data analysis. J. Appl. Crystallogr. 45, 342–350 (2012).

47. D. Franke, D. I. Svergun, DAMMIF, a program for rapid ab-initio shape determination in small-angle scattering. J. Appl. Crystallogr. 42, 342–346 (2009).

48. V. V. Volkov, D. I. Svergun, Uniqueness of shape determination in small-angle scattering. J. Appl. Crystallogr. 36, 860–864 (2003).

49. W. Wriggers, P. Chacón, Using Situs for the registration of protein structures with low-resolution bead models from X-ray solution scattering. J. Appl. Crystallogr. 34, 773–776 (2001).

50. E. C. Meng et al., UCSF ChimeraX: Tools for structure building and analysis. Protein Sci. 32, e4792 (2023).

51. Y. Furuike, J. Abe, A. Mukaiyama, S. Akiyama, Accelerating in vitro studies on circadian clock systems using an automated sampling device. Biophys. Physicobiol. 13, 235–241 (2016).

52. Y. Furuike et al., Elucidation of master allostery essential for circadian clock oscillation in cyanobacteria. Sci. Adv. 8, eabm8990 (2022).

53. S. Kumar et al., TimeTree 5: An Expanded Resource for Species Divergence Times. Mol. Biol. Evol. 39, msac174 (2022).

54. D. Franke et al., ATSAS 2.8: a comprehensive data analysis suite for small-angle scattering from macromolecular solutions. J. Appl. Crystallogr. 50, 1212–1225 (2017).

