## Supplementary material for "Molecular Shape Evolution of Cyanobacterial Circadian Clock Protein KaiC": SI Appendix

**This PDF file includes:**

Figures S1 to S9
Table S1
SI References

**Figures**

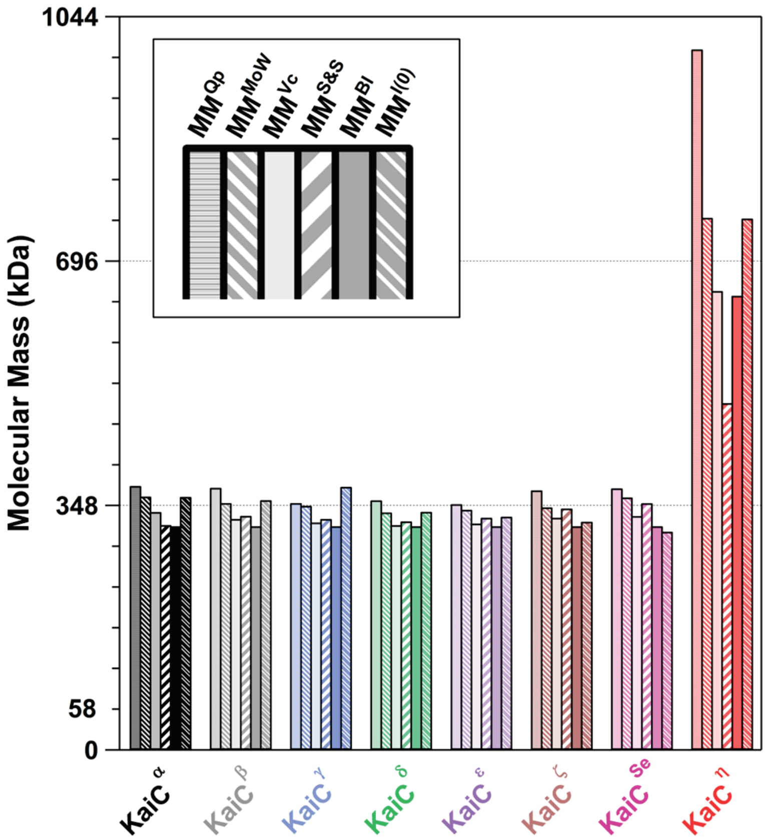


**Fig. S1.** Molecular masses of ancestral and modern KaiCs. The molecular masses (Table 1) were estimated based on apparent volume ($MM^{Qp}$) (1), Porod invariant ($MM^{MoW}$) (2), empirical volume of correlation ($MM^{Vc}$) (3), size and shape ($MM^{S\&S}$) (4), Bayesian inference ($MM^{BI}$) (1), and *I*(0) value of BSA standard ($MM^{I(0)}$) (5).

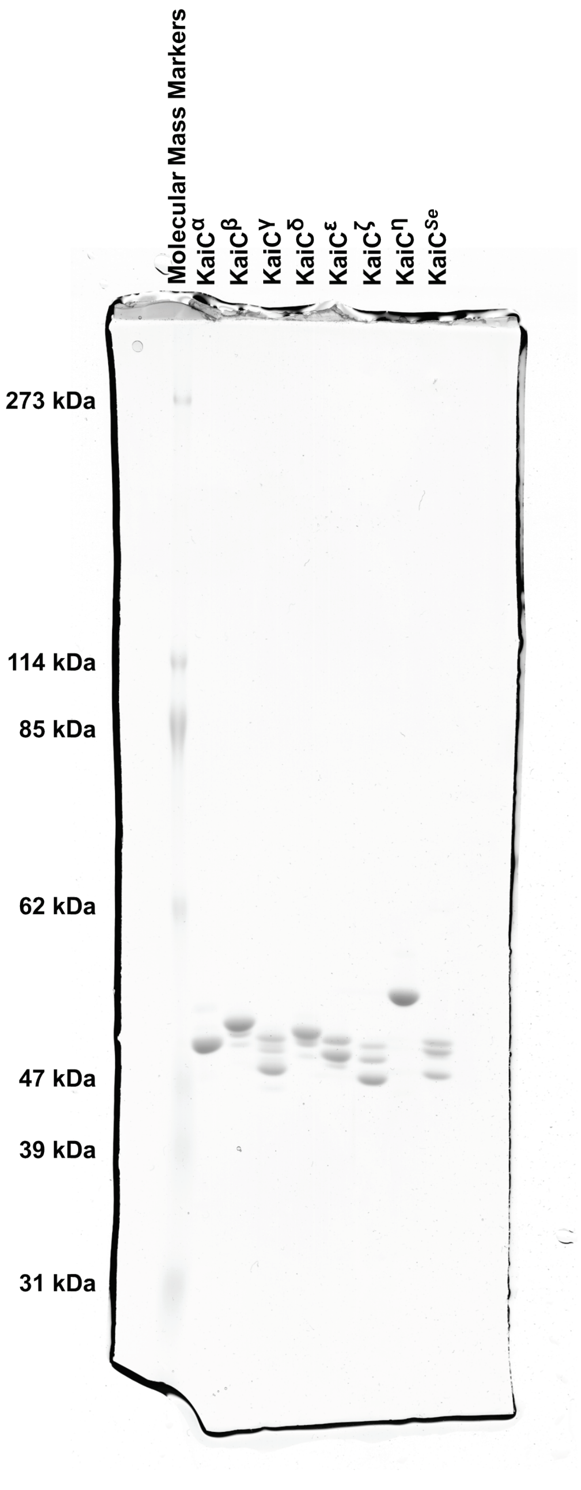


**Fig. S2.** Image of the original SDS-PAGE gel loaded with SAXS samples.

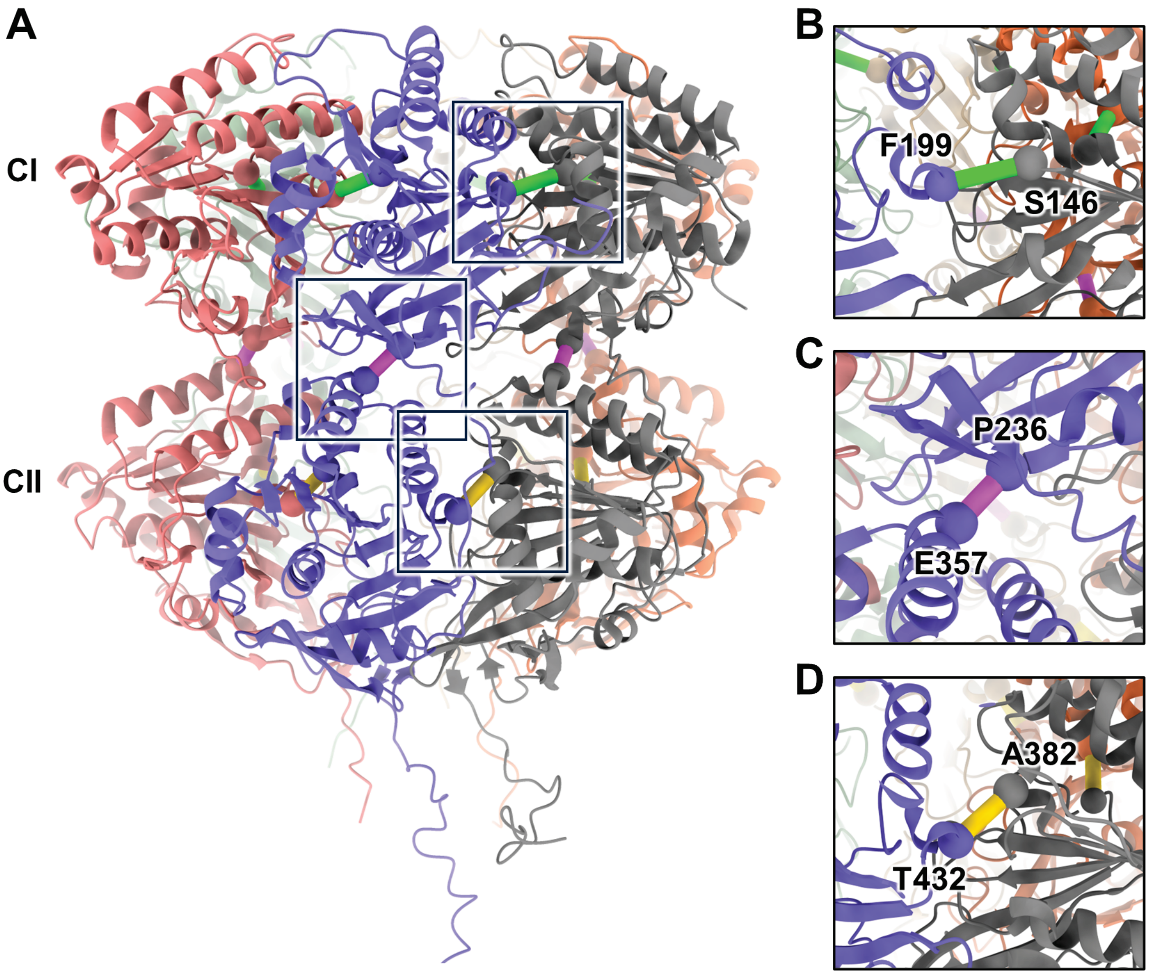


**Fig. S3.** Distance constraints during rigid-body refinements using CORAL (6). (**A**) Three kinds of soft distance constraints (green, magenta, and yellow: a penalty value of 10) mapped on the crystal structure of KaiC$^{Se}$ (PDB: 2GBL) (7). Relevant $C_\alpha$ atoms are drawn using sphere representation. (**B**) 9.5 Å for the $C_\alpha$ distance between F199 and S146 in the CI−CI interface. (**C**) 6.6 Å for P236 and E357 in the CI and CII domains, respectively. (**C**) 7.3 Å for T432 and A382 in the CII−CII interface.

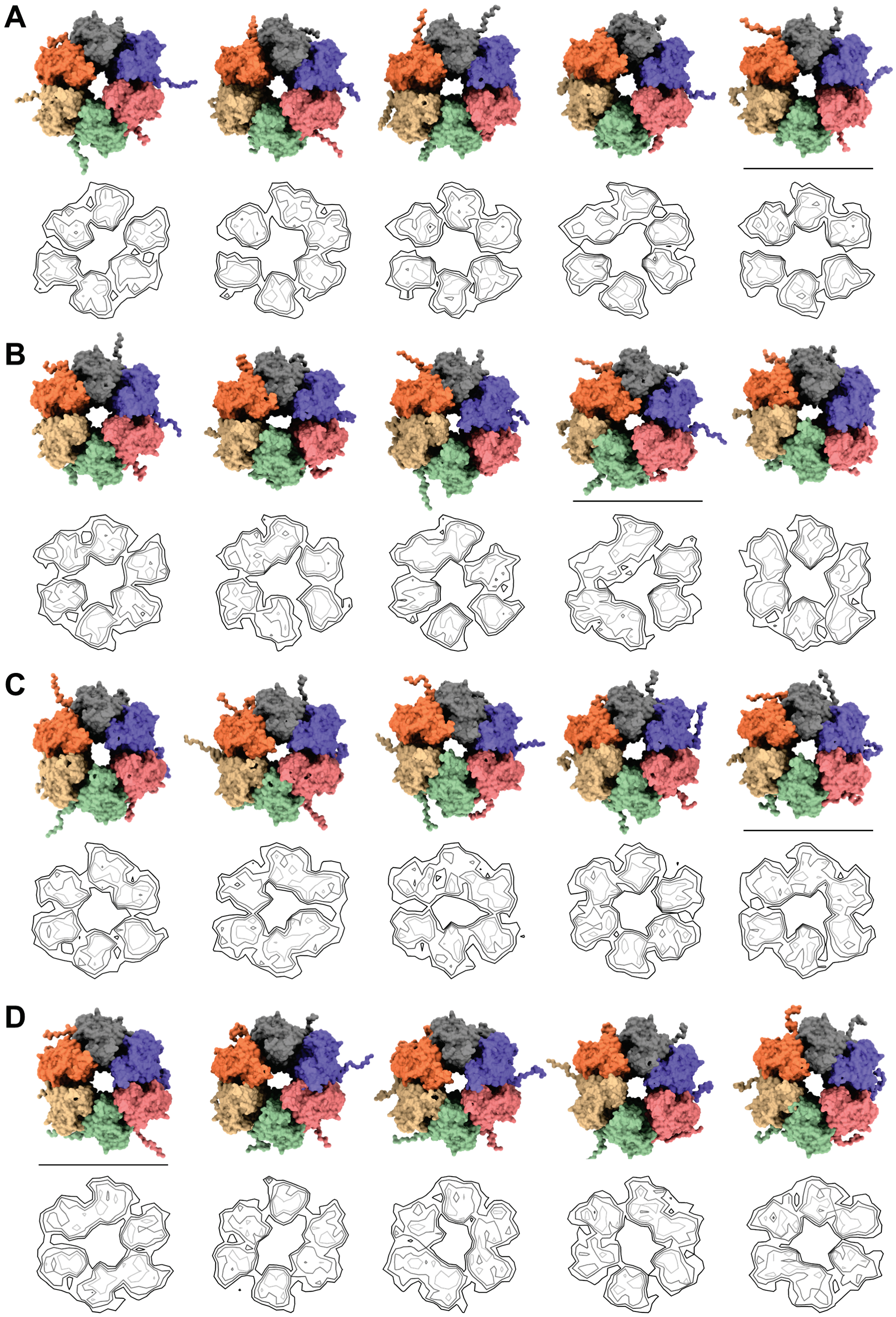


**Fig. S4.** Similarity and diversity of rigid-body models viewed from the CII-domain side. (**A**) KaiC$^{\alpha}$. (**B**) KaiC$^{\beta}$. (**C**) KaiC$^{\gamma}$. (**D**) KaiC$^{Se}$. C-terminal residues (from 501 to 519 or 525) are omitted for clarity of presentation. Contour images are the vertical projection (voxel spacing of 5 Å) of the CII ring (low: black, medium: gray, high: light gray). In the order from panel A to D, the black areas corresponding to the gaps between adjacent CII domains decrease. Underlined models correspond to the representative models shown in Fig. 4.

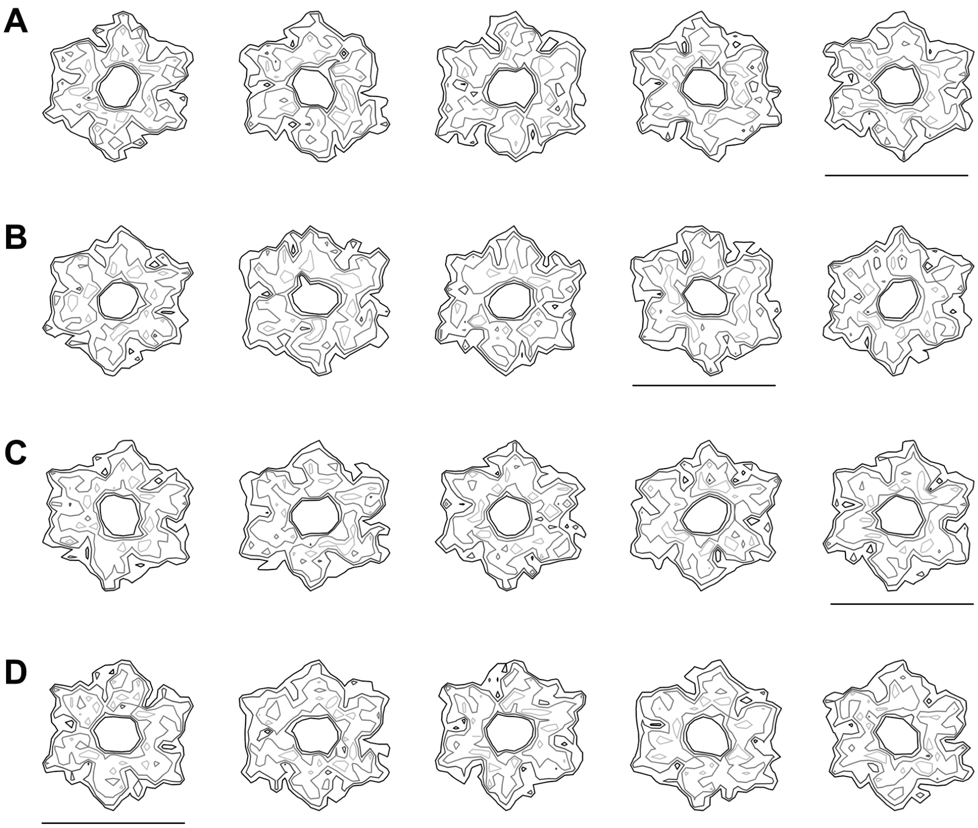


**Fig. S5.** Compact and near symmetric CI rings of rigid-body models. (**A**) KaiC$^{\alpha}$. (**B**) KaiC$^{\beta}$. (**C**) KaiC$^{\gamma}$. (**D**) KaiC$^{Se}$. Contour images are the vertical projection (voxel spacing of 5 Å) of the CI ring (low: black, medium: gray, high: light gray). N-terminal residues (from 1 to 13) are omitted for clarity of presentation. Underlined models correspond to the representative models shown in Fig. 4.

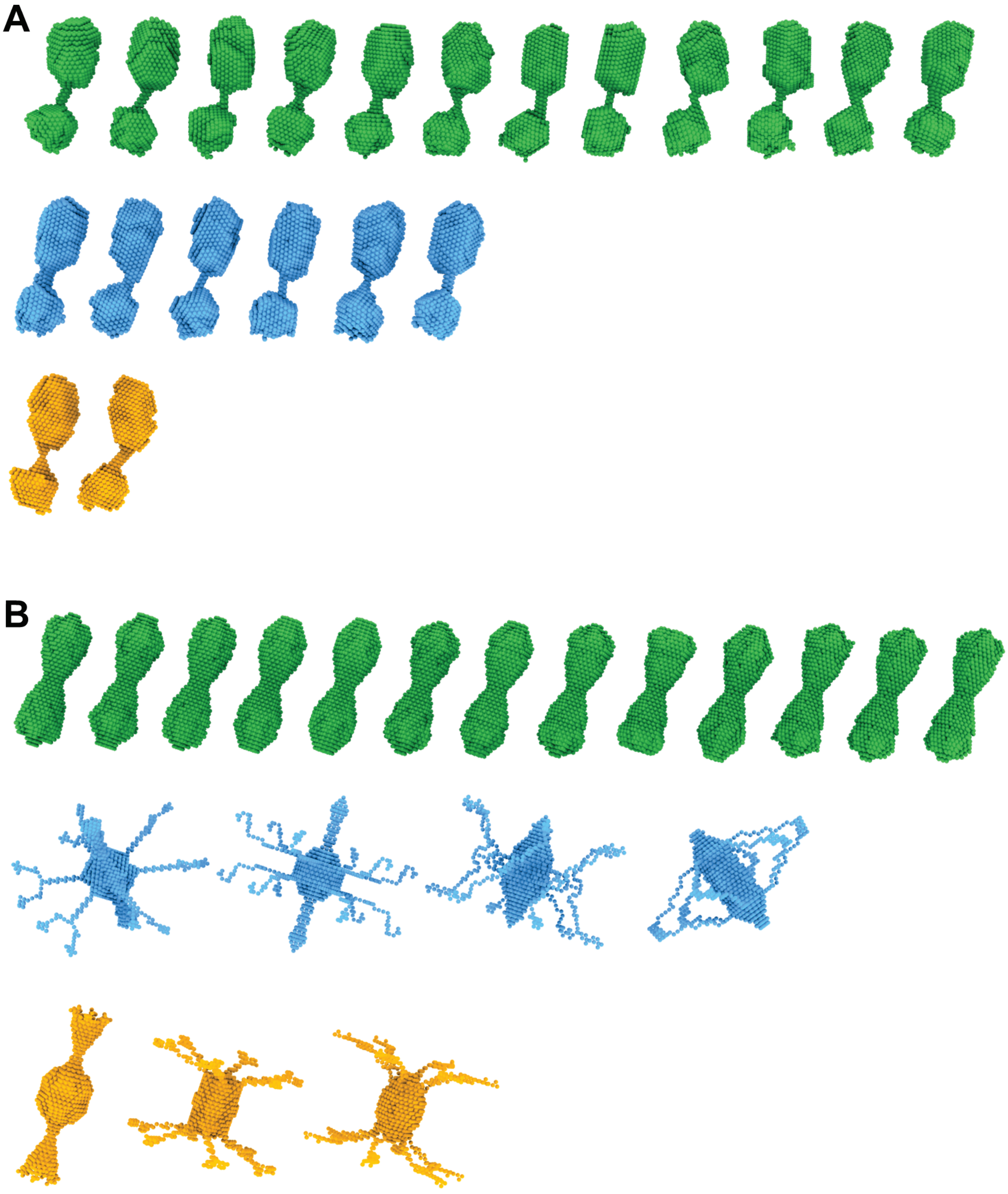


**Fig. S6.** Multiclass classification of *ab initio* models of KaiC$^{\eta}$ using DAMMAVER (8). (**A**) P1-symmetry reconstructions classified to one major (green) and two minor classes (cyan and orange). The low-resolution shape shown in Fig. 4E corresponds to the averaged model of the major class. (**B**) P62-symmetry reconstructions classified to one major (green) and two minor classes (cyan and orange). The low-resolution shape shown in Fig. 4F corresponds to the averaged model of the major class. See Table S1 for details on the refinement statics.

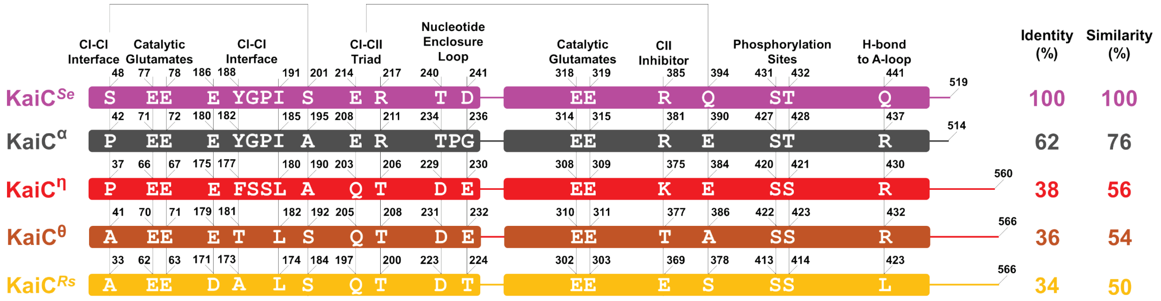


**Fig. S7.** Alignment of key residues and domain compositions of KaiC$^{Se}$, KaiC$^{\alpha}$, KaiC$^{\eta}$, KaiC$^{\theta}$, and KaiC$^{Rs}$. Sequence identities and similarities are given relative to KaiC$^{Se}$.

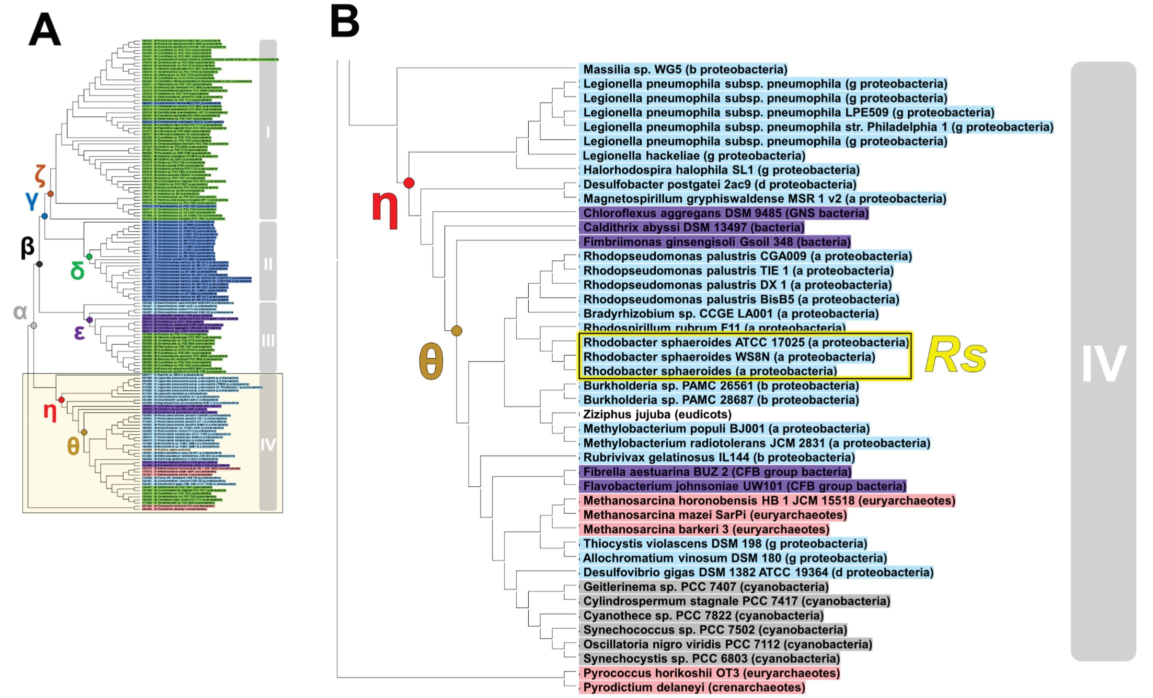


**Fig. S8.** Location of the θ node. (**A**) Overview and (**B**) expanded view of the phylogenic tree of KaiCs (9).

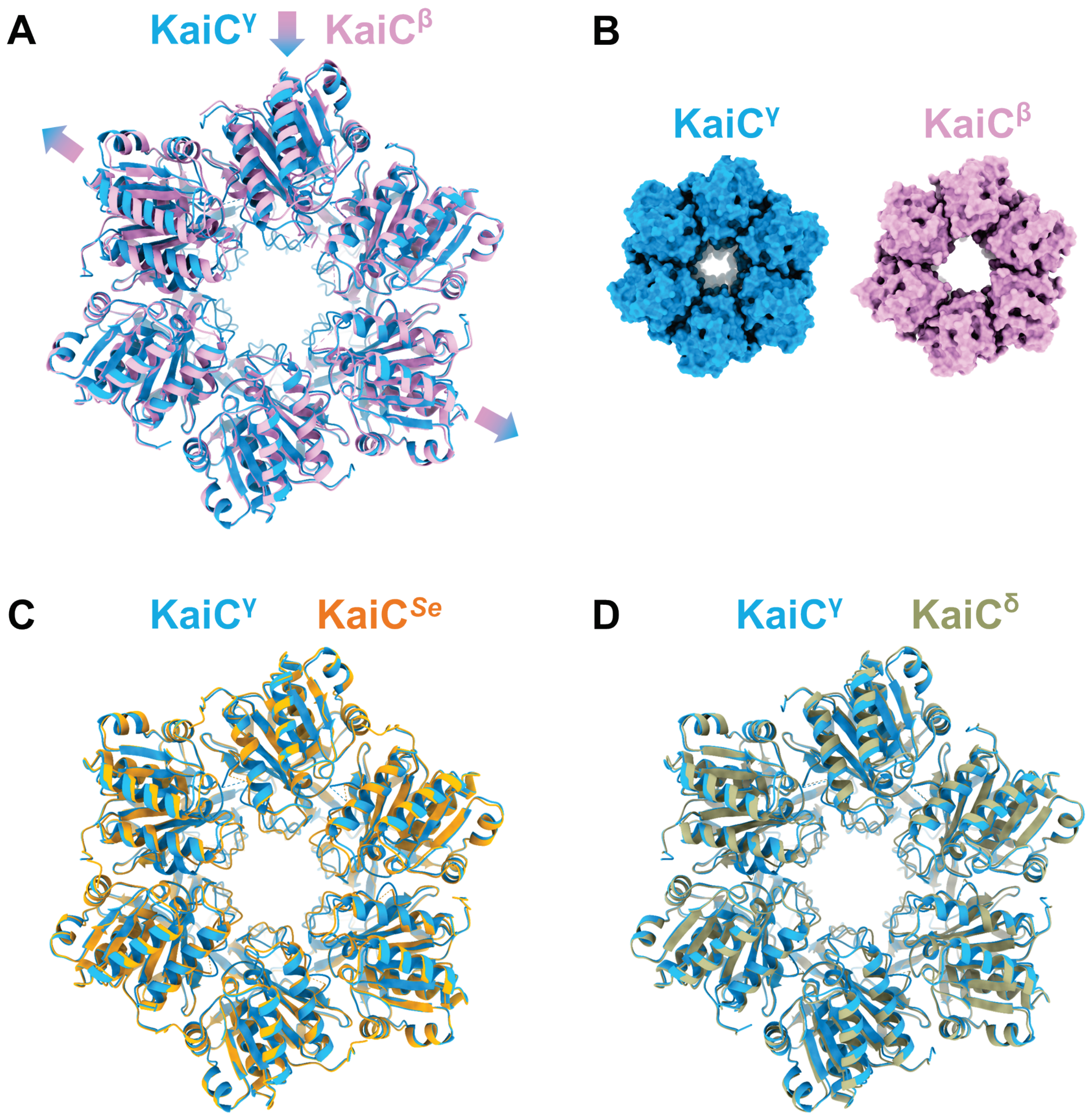


**Fig. S9.** Comparisons of compactness and symmetry of the CII rings in crystal structures. (**A**) KaiC$^{\gamma}$ (PDB: 8ZN6) and KaiC$^{\beta}$ (PDB: 8ZN5) (9). Arrows indicate the direction of changes from asymmetric (less compact) to symmetric (compact) CII rings. (**B**) Surface representations of KaiC$^{\gamma}$ and KaiC$^{\beta}$. (**C**) KaiC$^{\gamma}$ and KaiC$^{Se}$ (PDB: 7DXQ) (10). (**D**) KaiC$^{\gamma}$ and KaiC$^{\delta}$ (PDB: 8ZN7) (9).

### Tables

**Table S1.** SAXS-based modeling statistics for KaiC$^{\alpha}$, KaiC$^{\beta}$, KaiC$^{\gamma}$, KaiC$^{Se}$, and KaiC$^{\eta}$.

| | KaiC$^{\alpha}$ | KaiC$^{\beta}$ | KaiC$^{\gamma}$ | KaiC$^{Se}$ | KaiC$^{\eta}$ | |
|---|---|---|---|---|---|---|
| Modeling | Rigid-body<br>CORAL | Rigid-body<br>CORAL | Rigid-body<br>CORAL | Rigid-body<br>CORAL | *Ab initio*<br>DAMMIF | *Ab initio*<br>DAMMIF |
| Template | 2GBL | 2GBL | 2GBL | 2GBL | N.A. | N.A. |
| Symmetry | P1 | P1 | P1 | P1 | P1 | P62 |
| *Q*-range (Å$^{-1}$) | 0.01372−0.19409 | 0.01373−0.19411 | 0.01372−0.19409 | 0.01373−0.19413 | 0.00489−0.07942 | 0.00489−0.07942 |
| Total number of residues | 526 | 526 | 520 | 520 | N.A. | N.A. |
| Total number of dummy residues | 47 | 47 | 41 | 41 | N.A. | N.A. |
| Dummy residues | 1−13, 247−255,<br>501−525 | 1−13, 247−255,<br>501−525 | 1−13, 247−255,<br>501−519 | 1−13, 247−255,<br>501−519 | N.A. | N.A. |
| Total number of known residues | 479 | 479 | 479 | 479 | N.A. | N.A. |
| Known residues as rigid bodies | 14−246, 255−500 | 14−246, 255−500 | 14−246, 255−500 | 14−246, 255−500 | N.A. | N.A. |
| Constraints between $C_{\alpha}$ atoms | F199/S146: 9.5 Å<br>T432/A382: 7.3 Å<br>P236/E357: 6.6 Å | F199/S146: 9.5 Å<br>T432/A382: 7.3 Å<br>P236/E357: 6.6 Å | F199/S146: 9.5 Å<br>T432/A382: 7.3 Å<br>P236/E357: 6.6 Å | F199/S146: 9.5 Å<br>T432/A382: 7.3 Å<br>P236/E357: 6.6 Å | N.A. | N.A. |
| Number of reconstructed models | 5 | 5 | 5 | 5 | 20<br>Major: 12<br>Minor: 6<br>Other: 2 | 20<br>Major: 13<br>Minor: 4<br>Other: 3 |
| SQRT($\chi^2$)<br>(mean ± S.D.) | 1.45 ± 0.08 | 1.40 ± 0.03 | 1.31 ± 0.06 | 1.21 ± 0.03 | Major: 1.09 ± 0.04<br>Minor[1]: 1.06 ± 0.01<br>Minor[2]: 1.07, 1.05 | Major: 2.45 ± 0.62<br>Minor[1]: 122 ± 125<br>Minor[2]: 38.3 ± 13.5 |
| DAMAVER ICP[a]<br>(mean ± S.D.) | 5.67 ± 0.56 | 5.98 ± 0.73 | 4.49 ± 0.57 | 4.71 ± 0.52 | Major: 28.3 ± 1.8<br>Minor[1]: 27.1 ± 2.5<br>Minor[2]: 24.8 | Major: 5.56 ± 2.95<br>Minor[1]: 359 ± 401<br>Minor[2]: 184 ± 61 |

[a]ICP, iterative closest point.

**SI References**


1. N. R. Hajizadeh, D. Franke, C. M. Jeffries, D. I. Svergun, Consensus Bayesian assessment of protein molecular mass from solution X-ray scattering data. *Sci. Rep.* **8**, 7204 (2018).
2. H. Fischer, M. D. Neto, H. B. Napolitano, I. Polikarpov, A. F. Craievich, Determination of the molecular weight of proteins in solution from a single small-angle X-ray scattering measurement on a relative scale. *J. Appl. Crystallogr.* **43**, 101-109 (2010).
3. R. P. Rambo, J. A. Tainer, Accurate assessment of mass, models and resolution by small-angle scattering. *Nature* **496**, 477-481 (2013).
4. D. Franke, C. M. Jeffries, D. I. Svergun, Machine Learning Methods for X-Ray Scattering Data Analysis from Biomacromolecular Solutions. *Biophys. J.* **114**, 2485-2492 (2018).
5. S. Akiyama, Quality control of protein standards for molecular mass determinations by small-angle X-ray scattering. *J. Appl. Crystallogr.* **43**, 237-243 (2010).
6. M. V. Petoukhov *et al.*, New developments in the program package for small-angle scattering data analysis. *J. Appl. Crystallogr.* **45**, 342-350 (2012).
7. R. Pattanayek *et al.*, Analysis of KaiA-KaiC protein interactions in the cyano-bacterial circadian clock using hybrid structural methods. *EMBO J.* **25**, 2017-2028 (2006).
8. V. V. Volkov, D. I. Svergun, Uniqueness of shape determination in small-angle scattering. *J. Appl. Crystallogr.* **36**, 860-864 (2003).
9. A. Mukaiyama *et al.*, Evolutionary origins of self-sustained Kai protein circadian oscillators in cyanobacteria. *Nat. Commun.* **16**, 4541 (2025).
10. Y. Furuike *et al.*, Elucidation of master allostery essential for circadian clock oscillation in cyanobacteria. *Sci. Adv.* **8**, eabm8990 (2022).
